# Galectin-9 regulates dendritic cell polarity and uropod contraction by modulating RhoA activity

**DOI:** 10.1101/2023.10.30.564706

**Authors:** Guus Franken, Jorge Cuenca-Escalona, Isabel Stehle, Vince van Reijmersdal, Andrea Rodgers Furones, Rohit Gokhale, René Classens, Stefania Di Blasio, Yusuf Dolen, Annemiek B. van Spriel, Laia Querol Cano

## Abstract

Adaptive immunity relies on dendritic cell (DC) migration to transport antigens from tissues to lymph nodes. Galectins, a family of β-galactoside-binding proteins, control cell membrane organisation, exerting crucial roles in multiple physiological processes.

Here, we report a novel mechanism underlying cell polarity and uropod retraction. We demonstrate that galectin-9 regulates chemokine-driven and basal DC migration both in humans and mice, indicating a conserved function for this lectin. We identified the underlying mechanism, namely a deficiency in cell rear contractility mediated by galectin-9 interaction with CD44 that in turn regulates RhoA activity. Analysis of DC motility in the 3D tumour-microenvironment revealed galectin-9 is also required for DC infiltration. Moreover, exogenous galectin-9 rescued the motility of tumour-immunocompromised human blood DCs, validating the physiological relevance of galectin-9 in DC migration and underscoring its implications for DC-based immunotherapies.

Our results identify galectin-9 as a necessary mechanistic component for DC motility and highlight a novel role for the lectin in regulating cell polarity and contractility.

## Introduction

Dendritic cells (DCs) are the most potent antigen-presenting cell type, paramount for the induction of immune responses against pathogens and tumour cells. Cell migration endows DCs with the capacity to patrol their environment as well as to circulate between peripheral tissue and lymphoid organs, thereby linking innate and adaptive responses (Banchereau and Steinman, 1998; Delgado and Lennon-Dumenil, 2022). DC antigen uptake and maturation triggers the upregulation of specific surface proteins such as the chemokine receptor CCR7 that enables CCL19/CCL21 directed chemotaxis to secondary lymphoid organs and co-stimulatory molecules required for proper T cell activation (Acton et al., 2012; Randolph et al., 2005; Wculek et al., 2020). Rapid DC motility is crucial for their functions, which occurs in a so-called amoeboid manner, independent of integrins and mainly promoted by the existence of a front-rear polarity and actomyosin contractility (Lammermann et al., 2008). Amoeboid migration is mediated by the Rho family of small GTPases, key regulators of cytoskeletal dynamics that generate a polarised and dynamic activity balance at the front and rear of the cell (Hind et al., 2016; Sanchez-Madrid and Serrador, 2009). Illustrating this, Rac1 and Cdc42 located at the front of the cell drive filamentous actin polymerisation and directionality respectively, while RhoA at the cell rear, also called the uropod, mediates actomyosin contractility through Rho-associated kinase (ROCK) (Lammermann et al., 2009; Sanchez-Madrid and Serrador, 2009; Wong et al., 2006). In addition, polarised migrating DCs show enrichment of adhesion molecules such as CD44 at the uropod, which link the actin cytoskeleton to the cell membrane, promoting actin polymerisation in a RhoA-dependent manner (Bourguignon et al., 2003; Zhang et al., 2014b). Although DC migration dynamics are well characterised, the specific crosstalk between cell membrane events and intracellular cytoskeletal rearrangements that enable cell polarisation and underlie amoeboid migration remain unresolved.

CD44 is a highly glycosylated single-chain transmembrane receptor with crucial roles in cell adhesion and migration (Senbanjo and Chellaiah, 2017). Intracellularly, the cytoplasmic tail of CD44 interacts with RhoA, ERM (ezrin/radixin/moesin) or ankyrin to modulate cytoskeletal activation in response to extracellular cues (Bourguignon, 2008). Although the signalling events that control CD44-dependent cytoskeletal rearrangements are well-defined (Skandalis, 2023), the molecular mechanisms that regulate CD44 membrane distribution and how that influences cell migration remain elusive. Interestingly, galectins are required for CD44 nanoclustering and endocytosis at the plasma membrane of epithelial cells, suggesting galectin-mediated interactions are relevant for its spatiotemporal membrane organisation (Lakshminarayan et al., 2014).

Galectins are a group of lectins that display a conserved affinity for β-galactoside modifications on extracellular proteins and lipids (Johannes et al., 2018). All galectins contain one or two carbohydrate recognition domains that allow them to simultaneously interact with various glycosylated binding partners, thereby modulating the clustering, expression and activity of a large range of cell surface proteoglycans (Johannes et al., 2018; Liu and Stowell, 2023; Querol Cano et al., 2023). In addition to their extracellular functions, many galectins are also found in the cytosol (Hong et al., 2021; Johannes et al., 2018; Liu et al., 2002; Santalla Mendez et al., 2023) and in the nucleus where they participate in mRNA splicing (Coppin et al., 2017).

Galectin-9, encoded by the *Lgals9* gene, is a ubiquitously expressed tandem-repeat galectin, known to exert numerous roles in cancer, infection and inflammation (John and Mishra, 2016; Leffler et al., 2002; Tureci et al., 1997). Galectin-9 mediated functions are cell type dependent and are dictated by the spatiotemporal expression of its binding partners. Illustrating its versatility, galectin-9 was first characterised as an eosinophil chemo-attractant (Matsumoto et al., 1998), induces cell death and immune tolerance by binding T cell immunoglobulin-3 in T helper 1 (T_H_1) cells (Zhu et al., 2005), promotes the expansion of immunosuppressive macrophages (Arikawa et al., 2010) and monocytic myeloid-derived suppressor cells (Dardalhon et al., 2010) and negatively regulates B cell receptor signalling (Cao et al., 2018; Giovannone et al., 2018). Contrary to these reports implying an immunosuppressive role for galectin-9, we and others have previously identified galectin-9 to positively regulate DC biology (Dai et al., 2005; Li et al., 2023; Querol Cano et al., 2019; Santalla Mendez et al., 2023; Suszczyk et al., 2023). Nonetheless, the involvement of galectin-9 in immune cell migration has been insufficiently studied. For instance, dengue virus-infected DCs up-regulate galectin-9 expression and their ability to migrate towards CCL19 but whether galetin-9 is relevant for migration in non-pathological naïve DCs has not been addressed (Hsu et al., 2015). Furthermore, the mechanism(s) by which galectin-9 shapes cell motility are not delineated, with very few publications providing compelling evidence of how endogenous lectins modulate cytoskeleton rearrangements that underlie cell migration.

Here, we show galectin-9 is required for chemokine-driven and basal DC migration *in vitro* and *in vivo*, indicating an evolutionary conserved function for this lectin. We identified a defect in uropod retraction upon galectin-9 depletion, leading to a reduction in the fraction of active RhoA and its downstream signalling as the underlying mechanism. Importantly, we characterised the functional interaction between CD44 and galectin-9 at the plasma membrane of DCs as an essential driver of DC migration that integrates signals from external stimuli and dictates subsequent cytoskeletal rearrangements. Highlighting the physiological and translational relevance of our findings, exogenous galectin-9 was able to rescue the impaired migration capacity of tumour-immunocompromised human blood DCs, confirming the relevance of galectin-9 in DC motility.

## Results

### Galectin-9 is required for dendritic cell migration

Actin polymerisation regulates three-dimensional (3D) migration speed in DCs (Renkawitz et al., 2009) and our discovery that galectin-9 mediates actin contractility (Querol Cano et al., 2019) prompted us to study its involvement in governing DC migration. We investigated the functional consequence of galectin-9 depletion in DC migration using non-targeting siRNA-transfected and *LGALS9* siRNA-transfected monocyte-derived DCs (moDCs), herewith referred to as wild type (WT) and galectin-9 knockdown (galectin-9 KD) DCs respectively. Galectin-9 expression was almost completely inhibited (75-90 % reduction, Figures S1A and S1B) while analysis of HLA-DR, CD80, CD83 and CD86 surface expression showed a similar upregulation upon stimulation demonstrating no impairment in DC maturation upon galectin-9 depletion (Figures S1C and S1D). We first analysed chemokine-driven migration towards CCL21 using a transwell chamber containing a collagen gel. CCL21 induced DC migration in both WT and galectin-9 KD DCs but this was significantly decreased in the latter for all donors analysed (Figures 1A and 1B). Concomitantly, DCs were trapped in the collagen gel upon galectin-9 depletion (Figures S2A and S2B) demonstrating an involvement of galectin-9 in chemokine-driven DC migration. Next, DCs were subjected to a CCL21 gradient in the absence of collagen and also in this setup galectin-9 depletion resulted in an impaired migration (Figures S2C and S2D). Importantly, this migration defect was not attributed to the aberrant expression of the chemokine receptor CCR7 as this was equal in WT and galectin-9 KD DCs (Figure S1E). Next, we studied the effect of galectin-9 in basal 3D migration using live cell imaging microscopy. As shown, the average migratory velocity, mean square displacement and Euclidean distance were significantly diminished in galectin-9 KD DCs (Figures 1C, 1D, 1E and 1F respectively). Individual cell tracks demonstrate the diminished motility of galectin-9 KD DCs away from their initial location within the 3D collagen matrix (Figure 1G). Additional 3D migration assays in the presence of melanoma tumour spheroids were performed to investigate how galectin-9 depletion alters DC function in a physiologically relevant setup (Figure 2A). Time-lapsed video microscopy analysis demonstrated that also in this context, galectin-9 depletion significantly hampered DC velocity by approximately 40 % (average speed of 4.1 µm/min in WT *versus* 2.4 µm/min in gal-9 KD DCs) (Figure 2B) as well as their mean square displacement and Euclidean distance (Figures 2C and 2D). This effect was not tumour cell line specific as 3D assays performed with another melanoma line (BLM) yielded similar results (Figures S3A and S3B). Concomitant with a diminished cell velocity, galectin-9 KD DCs were found in lower numbers in the collagen surrounding the tumour spheroid and displayed a decreased infiltration rate compared to their WT counterparts (Figure 2E). Interestingly, WT DCs enhanced their migratory capacity both upon maturation and when present in the vicinity of a tumour whereas this increase was only marginal in galectin-9 KD DCs, suggesting that galectin-9 has a broad impact on the capacity of DCs to migrate (Figures S3C-S3E).

**Figure 1.**
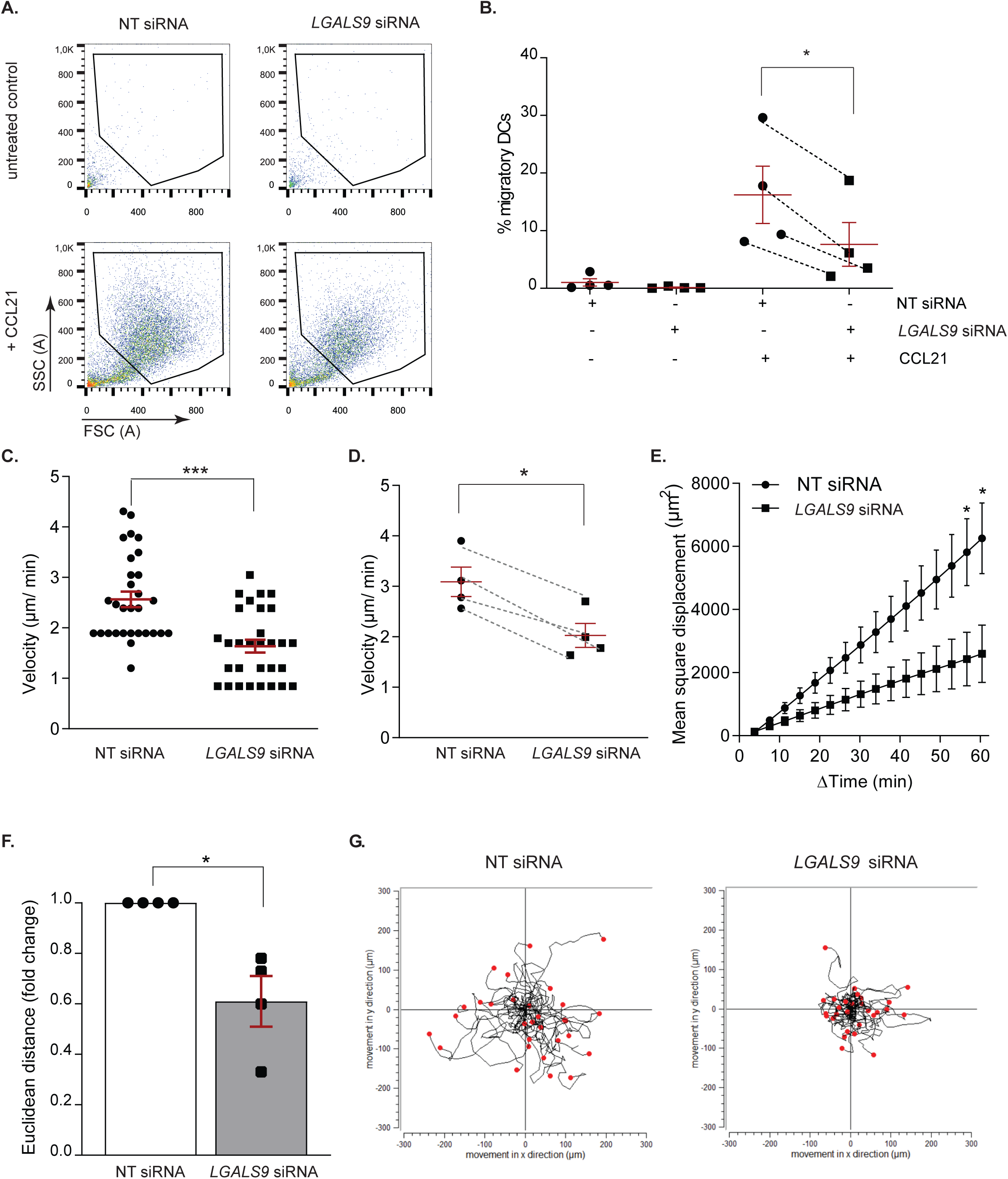
Galectin-9 depletion reduces dendritic cell migration. **A.** day 3 moDCs were transfected with either a *LGALS9* siRNA or a non-targeting (NT) siRNA as negative control, matured at day 6 for 48 h and subjected to a CCL21 chemokine migration assay using a transwell orverlaid with collagen matrix. Representative flow graphs from the migratory moDC fraction are shown. **B.** Quantification of donors shown in (A). Data shows the percentage of moDCs that migrated relative to input. Each symbol represents moDCs from one donor. **C.** moDCs were treated as per (A) and embedded within a 3D collagen matrix. The mean cell velocity of a representative donor is shown. **D – F.** Mean ± SEM cell velocity (D), mean square displacement (E) and Euclidean distance (F) of four independent donors. Twenty cells were analysed per donor and transfection. Lines in (D) connect matched NT and *LGALS9* siRNA transfected moDCs. Graph in (F) depicts relative Euclidean distance in *LGALS9* siRNA transfected moDCs after 60 minutes of tracking with respect to control cells. Graph depicts mean ± SEM of four independent donors. A one-way t-test was performed. **G.** Individual trajectory plots of NT and *LGALS9* siRNA transfected moDCs of one representative donor out of four analysed. End points of tracks are indicated by red dots. The black line indicates the overall movement in x and y direction (µm). Unpaired students t-test was conducted to compare NT siRNA and *LGALS9* siRNA transfected cells. * p < 0.05; *** p < 0.0001.

**Figure 2.**
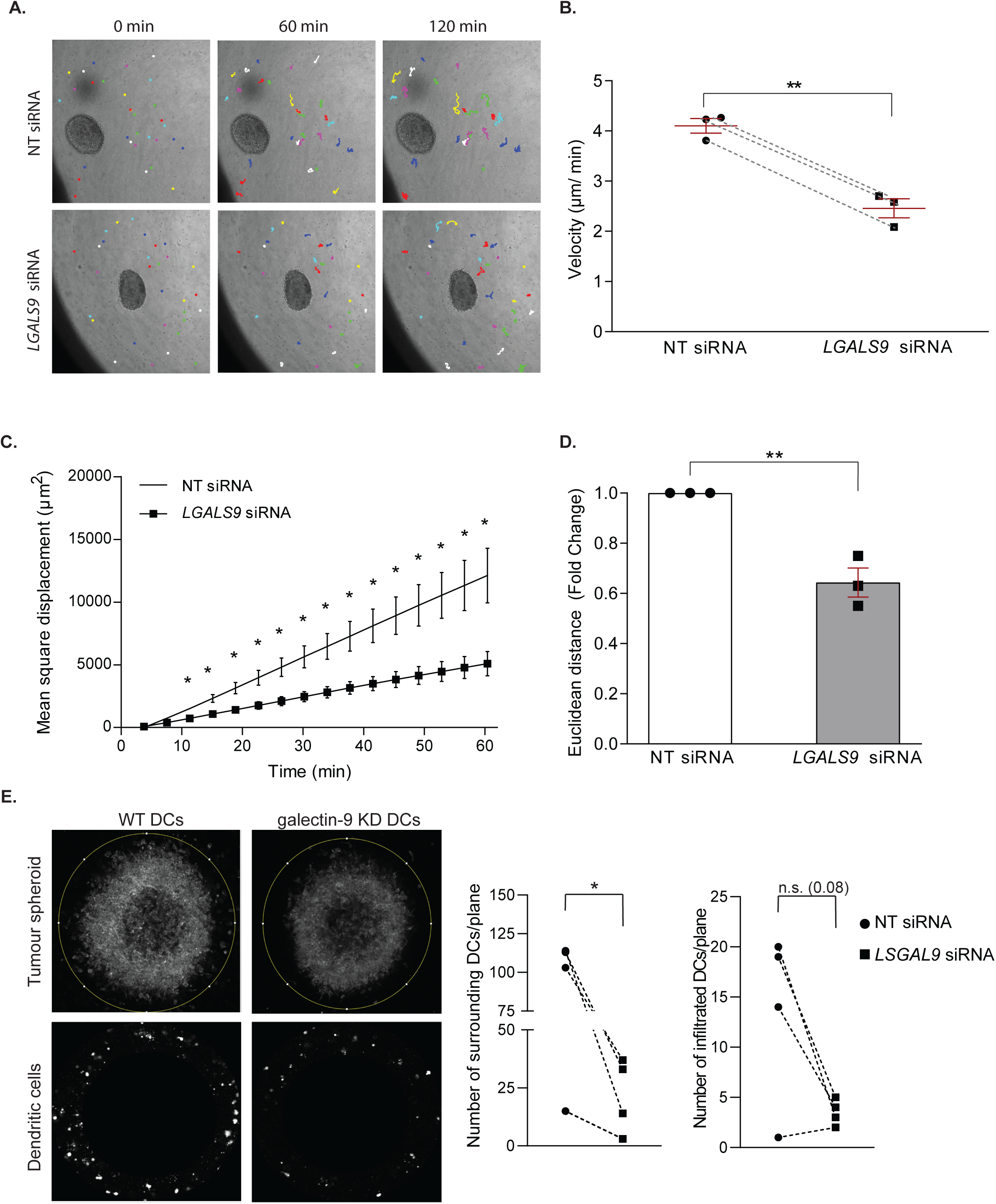
Migration of DCs within the tumour microenvironment is impaired upon galectin-9 depletion. **A.** Representative single cell tracking paths of NT or *LGALS9* siRNA moDC embedded in a 3D collagen matrix together with a Mel624 malignant melanoma spheroid. Dots indicate cell position at the specified time point whereas lines represent the tracking path covered by each cell from their initial position (time = 0 min). **B.** The cell velocity, the mean square displacement **(C)** and the Euclidean distance migrated after one hour **(D)** are depicted. Data represents mean value ± SEM of three independent donors. Euclidean distance is depicted as the mean value for galectin-9 depleted moDCs relative to the control group after 60 minutes of tracking. A one-way t-test was performed. In (B) lines connect matched NT and *LGALS9* siRNA transfected moDCs. **E.** Collagen matrices from (A) were fixed, stained for actin and the number of DCs surrounding (left graph) and infiltrated (right graph) in the spheroid calculated. Measurements were taken at the same tumour spheroid Z plane across conditions. Images are representative of the tumour spheroid and surrounding DCs. Graphs show mean ± SEM of three or four independent donors. Unpaired students t-test was performed between *LGALS9* and NT siRNA-transfected moDCs. n.s. = p > 0.05; * p < 0.05; ** p < 0.01.

Next, we sought to investigate whether galectin-9 function in DC migration was evolutionarily conserved using *in situ* migration assays on bone marrow-derived DCs (BMDC) from WT and *galectin-9* ^-/-^ (KO) mice (Figure 3A). WT and galectin-9 KO DCs were labelled with far-red or violet carboxyfluorescein succinimidyl ester (CFSE) dyes, mixed in equal numbers and co-injected into the same footpad or tail vein in host mice (Figure 3B). To rule out any involvement of galectin-9 present in the recipient animal both WT and galectin KO were employed as host mice. Donor DCs arriving in the draining popliteal and inguinal lymph nodes respectively were enumerated 48 h later via flow cytometry. The number of migratory *galectin-9*^-/-^ BMDCs was significantly reduced compared to WT DCs irrespective of the host genotype (galectin-9 WT or KO) (Figure 3C and 3D). Similar results were obtained in experiments in which violet-labelled WT and far-red labelled galectin-9 KO BMDCs were employed, ruling out any specific effect of the fluorescent dyes on migration (Figure 3C and 3D). CCL21 transwell chemotactic assays demonstrated an impairment in murine DC migration upon loss of galectin-9, confirming an evolutionary role for galectin-9 in driving DC motility (Figure S4).

**Figure 3.**
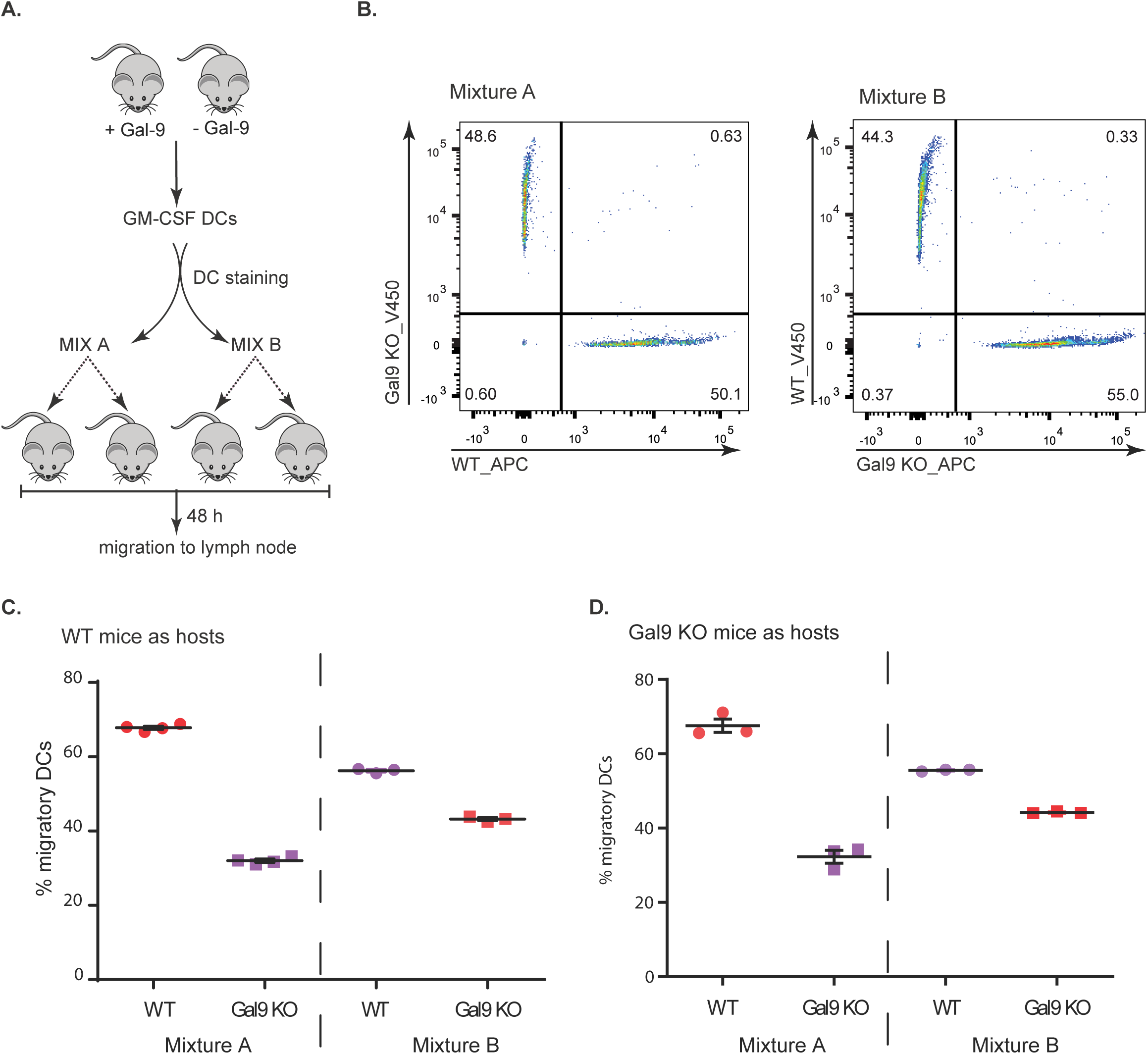
Galectin-9 function in dendritic cell migration is conserved *in vivo*. **A.** Scheme depicting experimental setup. WT and *galectin-9* ^-/-^ BMDCs were stained with far-red or violet CFSE dyes, mixed in equal numbers and injected into the footdpad or tail vein of donor mice. Forty eight hours later draining lymph nodes were isolated and donor BMDCs enumerated by flow cytometry. **B.** Representative flow cytometry plots depicting the cellular mixtures injected into recipient mice. In mixture A, WT BMDC were stained with far-red CFSE and galectin-9 ^-/-^ cells received violet CFSE. For mixture B, colours were interchanged to discard any effect of the dye in cell migration. **C** and **D.** Dendritic cell mixtures from (B) were injected into WT (C) and galectin-9 KO (D) host mice. Data depicts percentage of migratory donor BMDCs ± SEM. Each symbol represents values obtained for one draining lymph node.

Galectin-9 is located both in the cytosol as well as extracellularly (membrane-bound) in DCs (Querol Cano et al., 2019). To gain further insights into the molecular mechanisms underlying galectin-9 regulation of DC migration we cultured galectin-9 KD DCs with exogenous recombinant galectin-9 protein (gal-9 KD + rGal9 DCs). Analysis of galectin-9 expression revealed that exogenous protein restored surface-bound levels of galectin-9 while the cytosolic pool remained mostly depleted (Figure 4A). We next embedded WT, galectin-9 KD and gal-9 KD +rGal9 DCs in 3D collagen matrices to characterise their migration capacities. Restoring surface galectin-9 levels rescued the migration deficiency observed in galectin-9 KD DCs and no differences were observed in the velocity or the mean square displacement between WT and gal-9 KD + rGal9 DCs (Figures 4B and 4C, respectively). Moreover, individual cell tracks illustrate the ability of DCs to migrate upon restoring galectin-9 levels as galectin-9 KD + rGal9 DCs are indistinguishable from their WT counterparts (Figure 4D). Interestingly, the exogenous addition of galectin-9 rescued DC migration also after short incubations confirming the fundamental role of galectin-9 in driving DC migration (Figures S5A and S5B).

**Figure 4.**
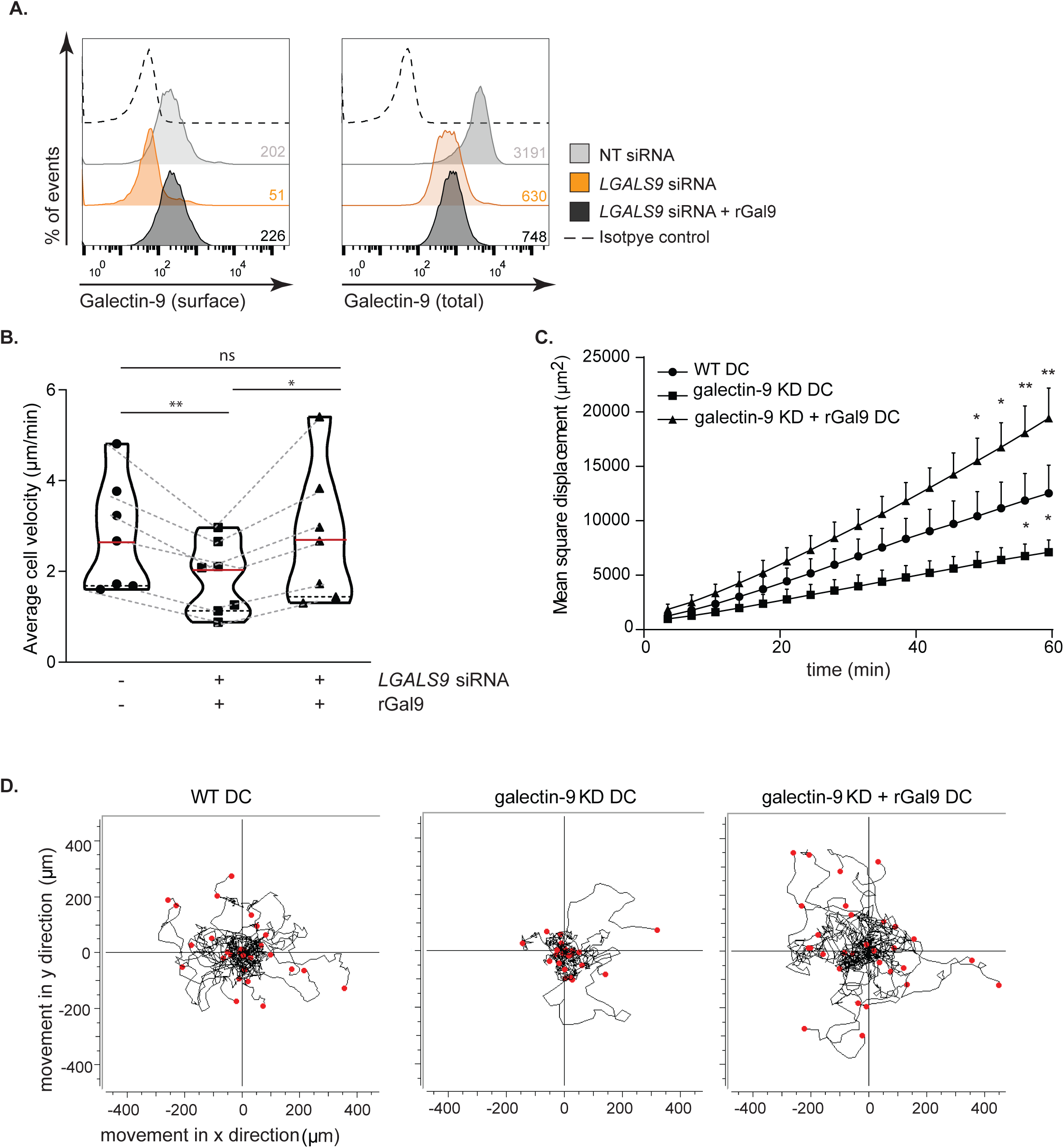
Dendritic cell migration relies on membrane-bound galectin-9 fraction. **A.** moDCs were transfected with *LGALS9* or a NT siRNA. Forty-eight hours after transfection, galectin-9 knockdown (galectin-9 KD) cells were treated with 1 µg/ml recombinant galectin-9 (rGal9) protein (gal-9 KD + rGal9) or nothing (WT) as negative control for 24-48 h. Surface only (left) and total (right) galectin-9 knockdown and treatment with exogenous protein were confirmed by flow cytometry 48 h after transfection. Grey population = WT; orange population = galectin-9 KD; black population = gal-9 KD + rGal9; dotted line = isotype control. Numbers in inset indicate geometrical mean fluorescence intensity (gMFI). **B** and **C.** moDCs treated as per (A) were embedded in 3D collagen matrices followed by live cell imaging to individually track cell migration. At least twenty cells were analysed for each donor and transfection or treatment. **B.** Violin plot showing average cell velocity of six individual donors. Each symbol represents one independent donor and lines connect paired donors. **C.** Mean square displacement was calculated and graph shows mean ± SEM of five individual donors. **D.** Individual trajectories of WT, galectin-9 KD and gal-9 KD + rGal9 moDCs. End points of tracks are indicated by red dots. Data shows representative donor out of six analysed. One-Way ANOVA test with a Bonferroni post-test correction was conducted between WT, galectin-9 KD and gal-9 KD + rGal9 moDC conditions. n.s. = p > 0.05; * = p < 0.05; ** = p < 0.01.

Overall, our results demonstrate that galectin-9 is required for both basal and chemokine-directed migration in DCs. Furthermore, this function appears to be evolutionarily conserved and is likely mediated by the surface-bound fraction of the lectin.

### Galectin-9 controls RhoA-mediated contractility in dendritic cells

To mechanistically resolve how galectin-9 dictates DC migration we morphologically characterised migratory WT, galectin-9 KD and gal-9 KD +rGal9 DCs in a 3D collagen matrix. WT cells contracted the uropod with concomitant forward movement (Figure 5A and Supplementary movie 1) whereas galectin-9 KD DCs were defective in their ability to contract the cell rear (Figures 5A, 5B and Supplementary movie 2). Remarkably, addition of exogenous galectin-9 protein rescued DC contractility and the uropod was not detected for abnormal lengths of time in gal-9 KD +rGal9 DCs (Figures 5A, 5B and Supplementary movie 3). Interestingly, DC elongation was not found to depend on galectin-9 expression (Figure 5C) implying that galectin-9 does not dictate cellular shape or length. Anterograde protrusion at the cell front is functionally dissociated from the retrograde contractility forces that mediate uropod retraction (Lammermann et al., 2008) and no differences were observed in the leading-edge protrusion formation suggesting galectin-9 is not involved in this process (data not shown).

**Figure 5.**
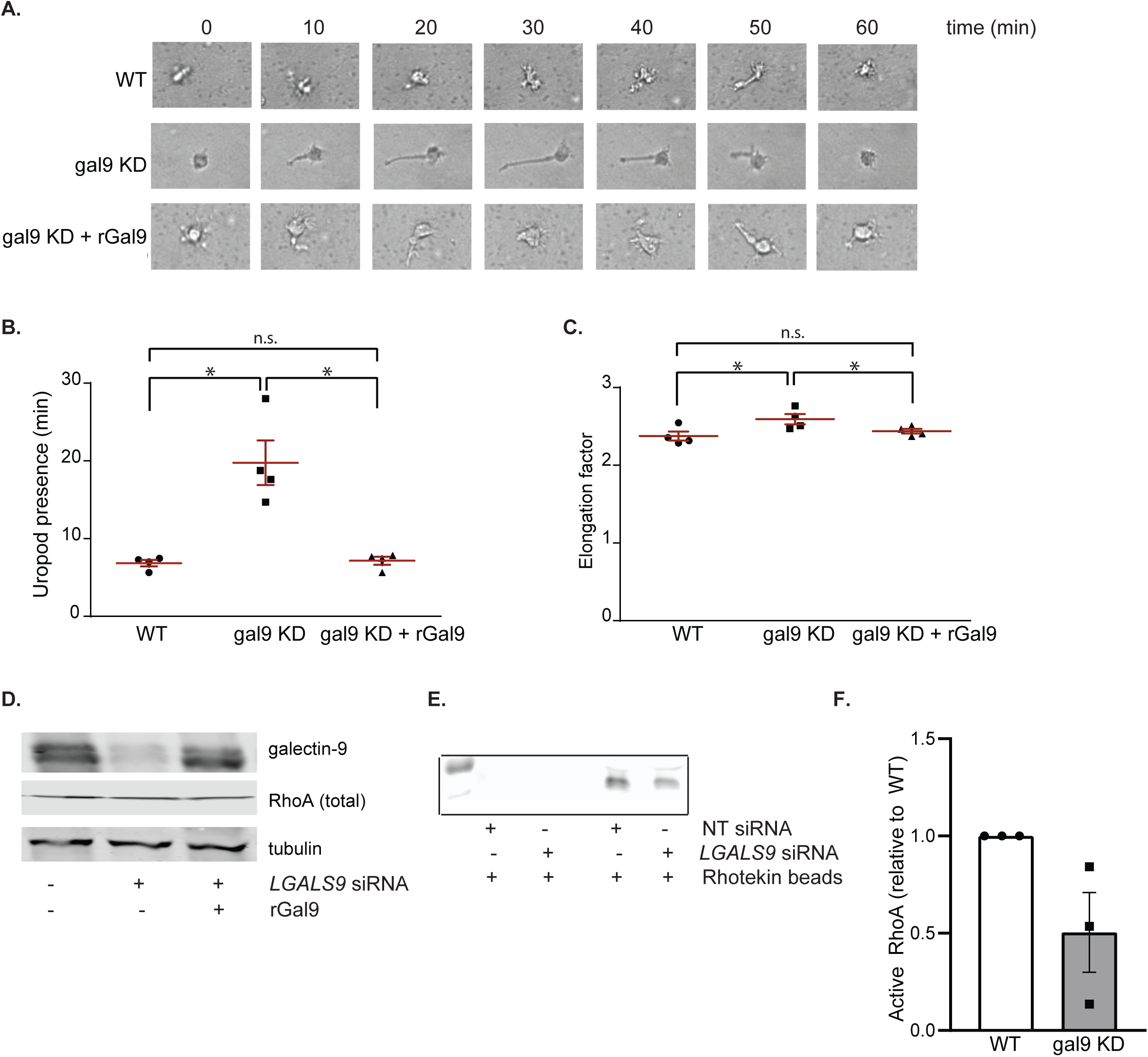
Galectin-9 regulates uropod contractility. **A.** Time-lapse sequence of a representative WT, galectin-9 depleted (gal9 KD) and gal-9 KD treated with recombinant galectin-9 protein (gal9 KD+rGal9) moDCs migrating in a 3D collagen gel. **B.** Duration of uropod presence (min) in WT DC, galectin-9 KD DC and gal-9 KD + rGal9 moDCs. **C.** Average cell elongation factor in WT DC, galectin-9 KD DC and gal-9 KD +rGal9 moDCs. Graphs depict mean ± SEM for four independent donors. Each symbol represents one independent donor. At least twenty cells were analysed for each donor and condition. One-Way ANOVA test with a Bonferroni post-test correction was conducted between WT DC, galectin-9 KD DC and gal-9 KD + rGal9 moDCs conditions. **D.** Total lysates from WT, galectin-9 KD and gal-9 KD +rGal9 moDCs were subjected to Western Blot and galectin-9 and total RhoA expression analysed. Tubulin was used as loading control. Immunoblot is representative of four independent experiments. **E.** Levels of active (GTP-bound) RhoA in WT DC and galectin-9 KD moDCs detected by immunoblotting. Rhotekin beads alone were used as negative control (lanes 2 and 3). **F.** Quantification of data shown in (E). Graph depicts relative active RhoA content in galectin-9 depleted DCs (gal9 KD) compared to relevant wild type (WT) control for three independent experiments. n.s. = p > 0.05, * = p < 0.05.

RhoA activity governs uropod contraction (Hind et al., 2016; Meili and Firtel, 2003) and thus we next examined if RhoA-mediated signalling was altered upon galectin-9 depletion. Although total RhoA levels did not differ across all conditions (Figure 5D), RhoA GTPase activity was markedly decreased in galectin-9 KD DCs (Figures 5E and 5F) in agreement with the aforementioned impairment in uropod retraction. Reverse phase protein array (RPPA) (Siwak et al., 2019) performed against more than 450 key functional proteins on WT, galectin-9 KD and gal-9 KD + rGal9 DCs whole lysates revealed a striking decline in either the expression or activity of proteins involved in cytoskeleton rearrangements in galectin-9 KD cells compared to WT or gal-9 KD + rGal9 DCs (Figure 6A). Remarkably, minimal differences in protein expression were detected between WT and gal-9 KD + rGal9 DCs indicating that treatment with exogenous galectin-9 protein rescues the DC signalling signature (Figure 6A). Enrichment pathway analysis (Zhou et al., 2019) performed on the differentially expressed proteins confirmed galectin-9 is a positive regulator of cell motility (Figure 6B). Validating our analysis, cytokine signalling was also found to be positively regulated in WT compared to galectin-9 KD DCs, which we have previously reported (Santalla Mendez et al., 2023). RPPA analysis revealed that the active form of P21 activated kinase 1 (PAK1) (PAK_Thr423), an activating Ser/Thr kinase downstream of RhoA, is downregulated in galectin-9 KD DCs whereas treatment with exogenous galectin-9 protein (gal-9 KD + rGal9 DCs) rescued its levels to those found in WT cells (Figure 6A). Total levels of PAK1 did not differ across conditions suggesting galectin-9 is not involved in regulating its expression. Analysis of phosphorylated and total PAK1 levels in multiple donors validated our RPPA data and confirmed PAK to be differentially activated in response to galectin-9 cellular levels (Figure 6C). Interestingly, we did not see any differences in other downstream RhoA signalling molecules (mDia and phosphomyosin light chain (PiMLC)) (Figures 6D and 6E) upon galectin-9 depletion or over-expression suggesting galectin-9 regulates RhoA-mediated contractility by mechanisms that are independent of MLC phosphorylation. To further study the link between galectin-9 and RhoA, we sought to investigate which genes positively correlate with *lgals9* expression and the pathways they mediate. We obtained a list of the top 50 genes correlating with *lgals9* using the ULI RNA-seq dataset (GSE109125) from the Immunological Genome Project (ImmGen) and performed a functional pathway enrichment analysis using the Reactome dataset (Griss et al., 2020) (Figure S6A). Data obtained showed a significant enrichment in both the Rho GTPase effectors and signalling by Rho GTPases pathways confirming a correlation between galectin-9 and RhoA activity (Figure S6B). Overall, our data shows that galectin-9 regulates cell rear contractility by modulating RhoA retrograde activity and Pak1-mediated downstream signalling.

**Figure 6.**
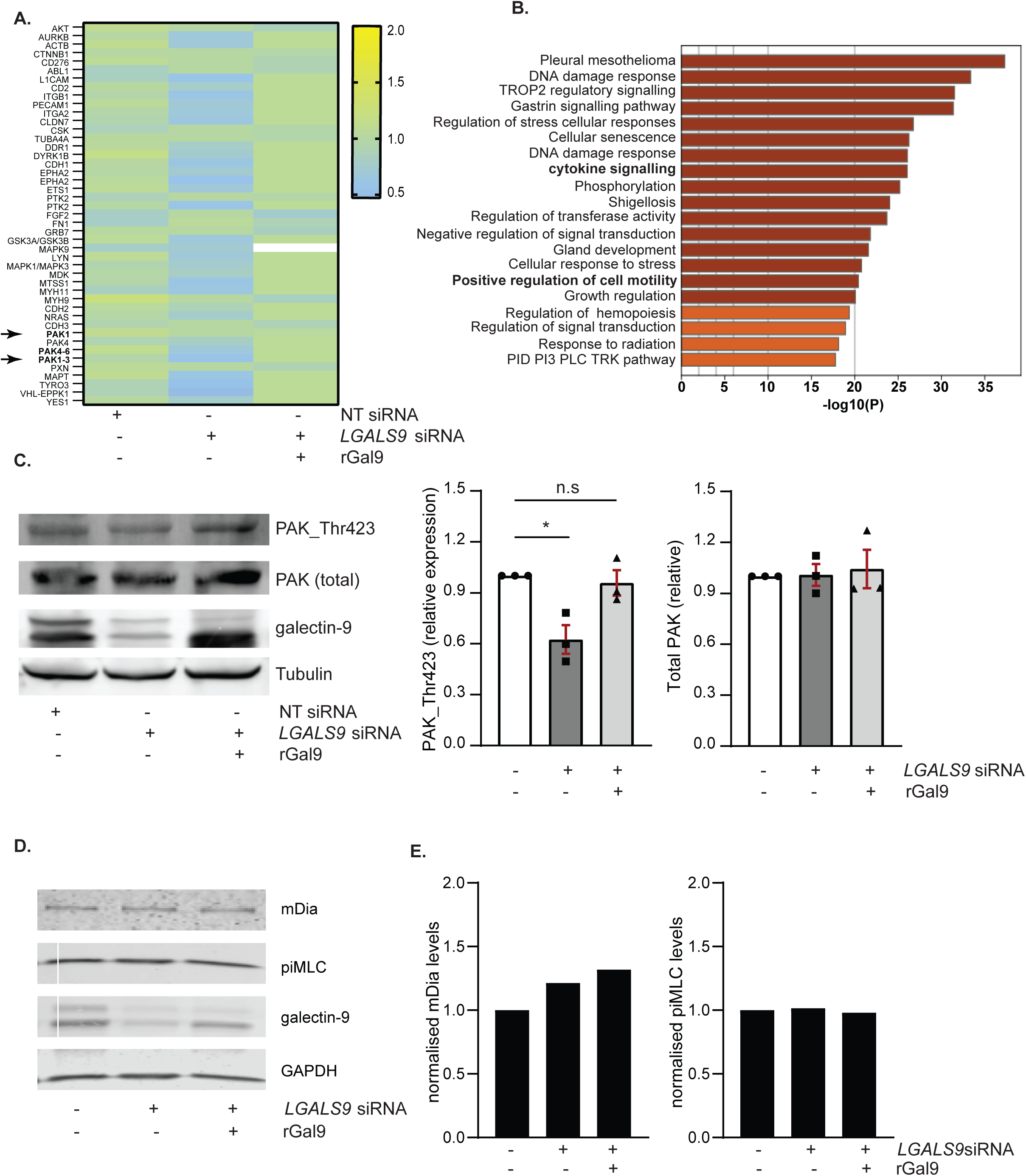
RhoA downstream signalling is impaired in galectin-9 depleted DCs. **A.** Heat map of the proteins involved in cytoskeletal rearrangements analysed by RPPA in WT, galectin-9 depleted (galectin-9 KD) and galectin-9 KD treated with recombinant protein. **B.** Metascape pathway enrichment analysis from all proteins found to be differentially regulated by RPPA between WT and galectin-9 KD moDCs. **C.** Total lysates from WT, galectin-9 KD and gal-9 KD + rGal9 moDCs were subjected to Western Blot and total PAK1, PAK_Thr423 and galectin-9 protein expression analysed. Tubulin was used as loading control. Immunoblot is representative of three independent experiments. Graph shows quantification of PAK1_Thr423 and total PAK1 content normalised to tubulin in each sample using ImageJ. Graph shows mean ± SEM of three independent donors. Two-way ANOVA followed by Dunnett’s test for multiple comparison was performed. n.s = p > 0.05; * = p < 0.05. **D.** Total lysates from WT, galectin-9 KD and gal-9 KD +rGal9 moDCs were subjected to Western Blot and mDia, phosphomyosin light chain (piMLC) and galectin-9 expression analysed. Tubulin was used as loading control. Immunoblot is representative of four independent experiments. **E.** Quantification of mDia and piMLC content shown in (D) normalised to tubulin in each sample using ImageJ.

The transmembrane adhesion glycoprotein CD44 interacts with RhoA via its cytoplasmic tail. We performed co-immunoprecipitation (IP) experiments using galectin-9 specific antibodies and identified CD44 to interact with galectin-9 in moDCS (Figure 7A). As positive control, Vamp-3 was also enriched in the galectin-9 IP compared to isotype control as previously reported (Santalla Mendez et al., 2023). To characterise the functional interaction between CD44 and galectin-9 in DCs we treated naive moDCs with a blocking antibody that prevents galectin-9 binding to CD44 (Iqbal et al., 2022). Interestingly, abrogating CD44-galectin-9 interaction markedly diminished DC migration in transwell chemotactic assays (Figure 7B). We next assessed the effects of blocking CD44-galectin-9 binding in 3D collagen assays as they pose a more physiological model. Blocking CD44 binding to galectin-9 also decreased cell motility although to a lesser extent than that obtained upon galectin-9 depletion (Figure 7C and 7D). Interestingly, pre-incubation of galectin-9 KD DCs with the anti-CD44 blocking antibody prior to the addition of recombinant galectin-9 protein abolished the rescue in cell motility observed upon restoring galectin-9 levels (Figure 7C and 7D). Importantly, galectin-9 depletion or overexpression did not affect antibody binding to CD44 (Figure 7E) or CD44 surface expression (Figure 7F). Overall, our data identifies the functional interaction between CD44 with galectin-9 at the surface of DCs as the molecular mechanism by which galectin-9 modulates RhoA activity, thereby regulating actin polymerisation and contractility.

**Figure 7.**
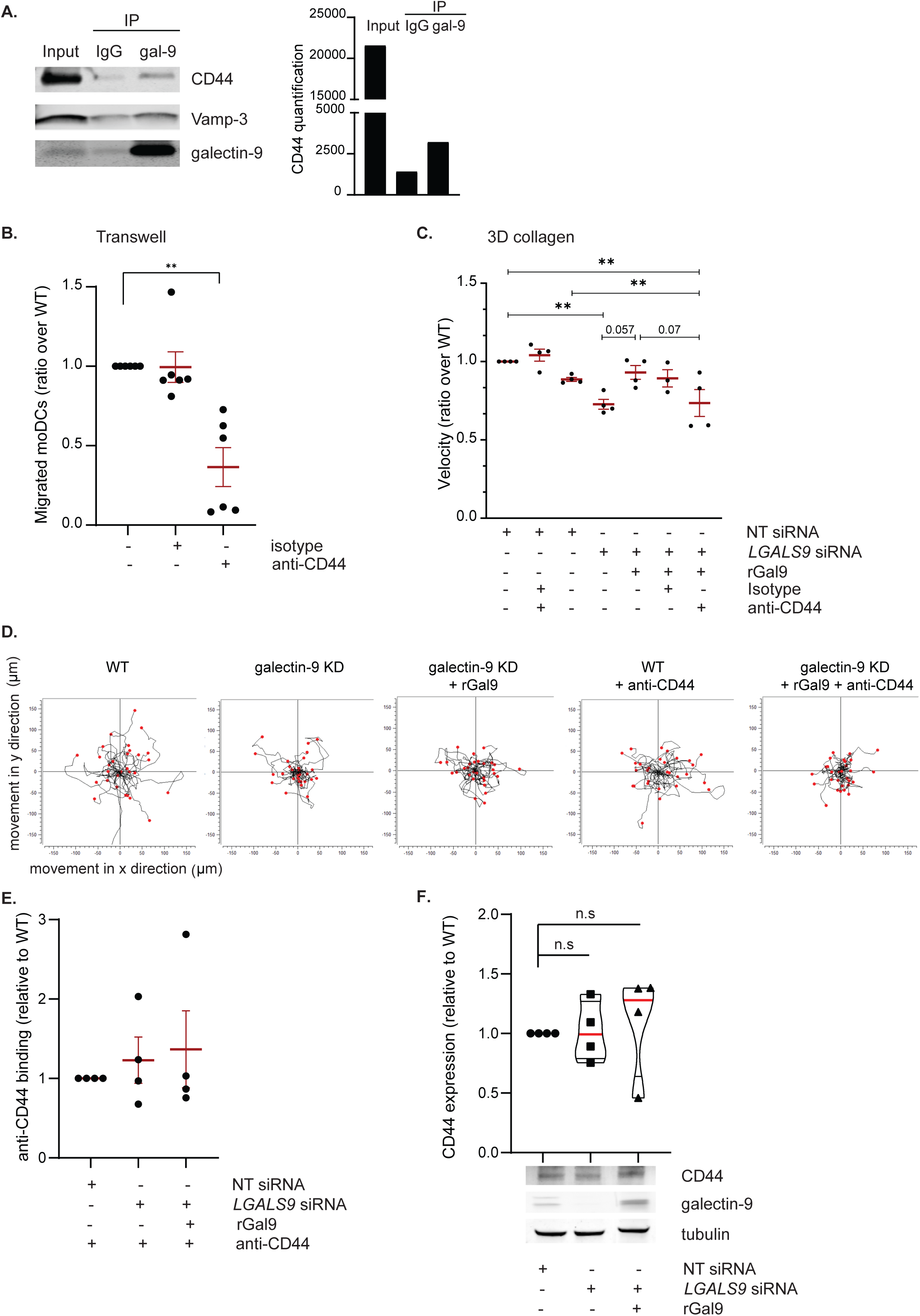
Galectin-9 interaction with CD44 regulates DC migration. **A.** moDCs were lysed and whole cell extracts incubated with either anti-galectin-9 specific antibody or isotype control. Immunoprecipitated complexes were resolved and probed with galectin-9, Vamp-3 and CD44 specific antibodies. Graph shows quantification of CD44 signal. **B.** moDCs were treated with 10 µg/ml of an anti-CD44 blocking antibody or isotype control for 30 min prior to being placed in the upper chamber of a transwell system and subjected to a CCL21 chemokine gradient for 3 h. Migrated cells were measured by flow cytometry. Graph shows mean percentage ± SEM of migratory DCs relative to total number of seeded cells for six individual donors. **C.** *LGALS9* siRNA or a non-targeting siRNA (NT) transfected moDCs were treated with 10 µg/ml of an anti-CD44 blocking antibody or isotype control for 30 min prior to being treated with 1 µg/ml recombinant galectin-9 protein (rGal9) or nothing as negative control for further 30 min. Cells were then embedded in 3D collagen matrices and individual cell tracking performed. Graph show mean ± SEM of four independent donors. At least twenty cells were analysed for each donor and transfection or treatment. **D.** Representative individual trajectory plots from data shown in (C). End points of tracks are indicated by red dots. The black line indicates the overall movement in x and y direction (µm). Cell genotype and treatments are indicated above each graph. **E.** WT, galectin-9 KD and galectin-9 KD + rGal9 moDCs were treated with anti-CD44 antibody and antibody binding measured by flow cytometry. Graph shows antibody binding for each donor relative to the corresponding WT signal. Data represents mean ± SEM for four independent donors. **F.** Total lysates from WT, galectin-9 KD and gal-9 KD + rGal9 moDCs were subjected to Western Blot and CD44 and galectin-9 expression analysed. Tubulin was used as loading control. Immunoblot is representative of four independent experiments. Violin plots depict mean CD44 content in galectin-9 KD and gal-9 KD + rGal9 moDCs normalised to tubulin and relative to the corresponding WT sample.

### Galectin-9 is sufficient to rescue migration in tumour-immunosuppressed primary DCs

To further validate the importance of galectin-9 on DC migration in a model that better recapitulates a physiological setup, the human blood DC subset conventional DCs type 2 (cDC2s) was used. Mature cDC2s were treated with melanoma-derived conditioned medium (CM) to impair their migratory capacity, after which galectin-9 was provided to assess its ability to rescue cDC2 migration (Figure 8A). Migratory capacity of DCs was determined in a transwell migration assay towards the chemokines CCL19 and CCL21 for three hours (Figure 8B). Exposure of mature cDC2s to melanoma-derived CM led to the downregulation of surface galectin-9 and CCR7 expression levels (Figure 8C and 8D). In line with the phenotype data, tumour-primed cDC2s exhibited a lower migratory capacity towards the chemokines CCL19 and CCL21 compared to untreated mature cDC2s (Figure 8E). Remarkably, addition of exogenous galectin-9 rescued cDC2 migration towards the chemokines CCL19 and CCL21 (Figure 8E). CCR7 expression levels remained unchanged during galectin-9 addition (Figure 8D and Supplementary Figure 7B), indicating this superior migratory capacity to be dependent on galectin-9 presence and not on an altered CCR7 expression. Taken together, these results validate the relevance of galectin-9 for the migration capacity of naturally occurring DC subsets in a tumour model and illustrate the therapeutic value of intervening galectin-9 signalling axis to restore the migration of tumour-immunocompromised DCs.

**Figure 8.**
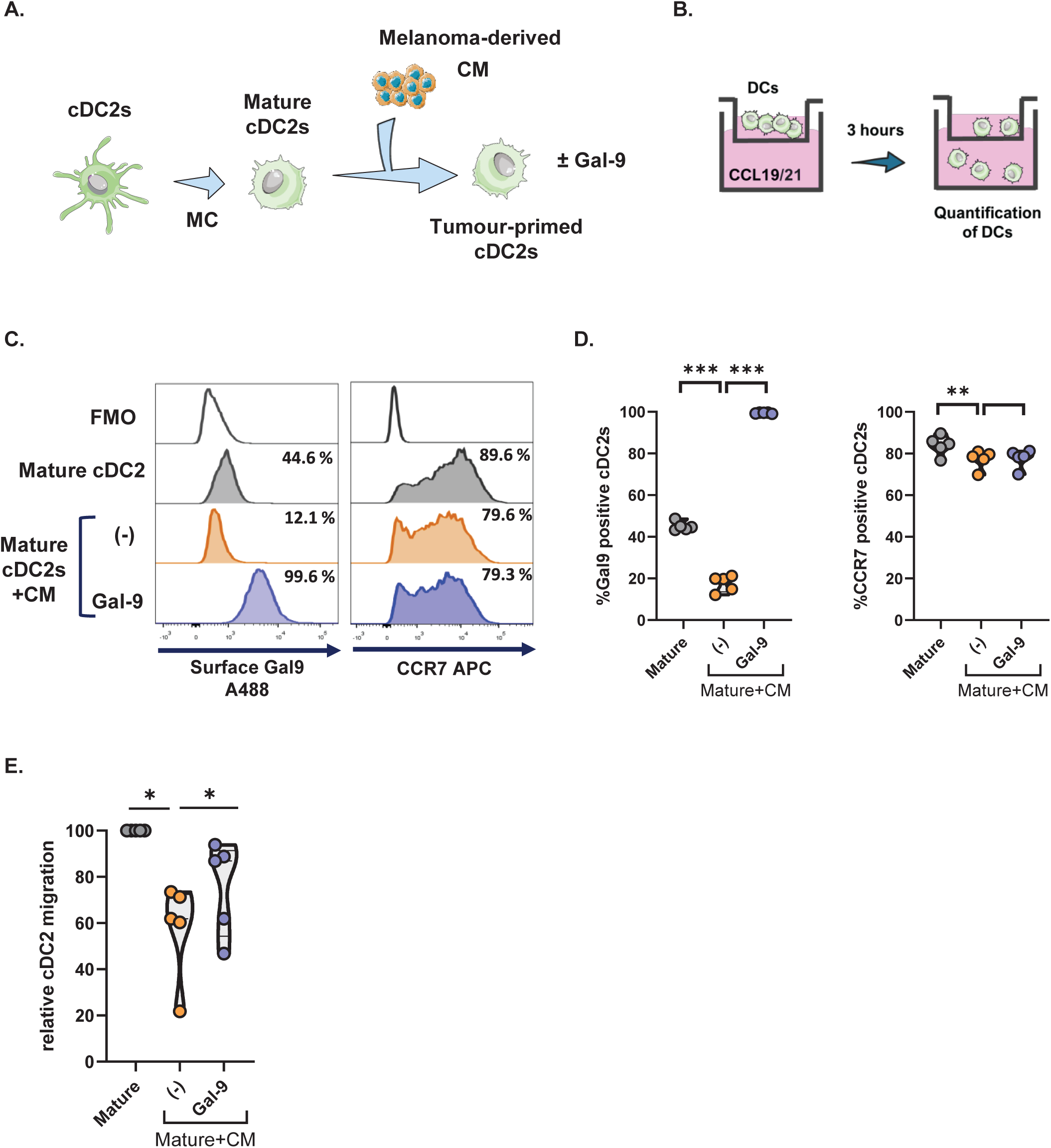
Galectin-9 restores chemokine-driven cDC2 migration. **A.** Schematic representation of the experimental setup. Primary human cDC2s were matured overnight with a maturation cocktail (MC) for 24 h prior to being harvested and replated in the presence of melanoma-derived conditioned medium (CM) for 24 h. Exogenous galectin-9 was supplemented for the last 2 h of cDC2 incubation with melanoma-derived CM. Tumour-primed cDC2s were collected and analysed for the surface expression levels of galectin-9 and CCR7 or for their migratory capacity. **B.** Schematic representation of the transwell migration assay. cDC2 were seeded in the top chamber of a transwell chamber containing a 5 µm porus membrane and subjected to a chemokine gradient of the CCR7-ligands CCL19 and CCL21. Migratory cDC2s were collected after 3 h and quantified. **C.** Histograms showing the surface expression of galectin-9 and CCR7 of a representative cDC2 donor analysed by flow cytometry. **D.** Percentage of positive cDC2s for galectin-9 and CCR7 for each of the indicated treatments. **E.** Relative cDC2 migration under each treatment determined by normalising each treatment to the migration given by mature cDC2s unexposed to the melanoma-derived CM for every donor. Violin plots in (D) and (E) show mean of five independent donors. One-way ANOVA followed by Dunnett’s test for multiple comparison was performed. * p < 0.05; ** p < 0.01; *** p < 0.001.

## Discussion

Single cell migration is an ubiquitous phenomenon in mammalian cell biology with cells mostly displaying either a mesenchymal or an amoeboid migration mode. The latter is employed by dendritic cells (DCs), allowing for a fast and autonomous migration, driven by actin polymerisation and actomyosin contractility forces. This is essential to rapidly shuttle antigens from peripheral tissues to lymphoid organs. However, how environmental cues and cell membrane organisation integrate and modulate cytoskeleton remodelling and contractility remains poorly characterised.

In this study, we uncovered a role for galectin-9 in DC migration and report a novel function in controlling rear actin contractility *via* its interaction with CD44 at the cell surface that in turn regulates RhoA cytosolic activity (Figure 9). Galectin-9 depletion significantly hampered basal and chemokine-driven migration in human and murine DCs, suggesting an evolutionary conserved function for the lectin. We also reveal that immunosuppressed blood cDC2 cells express low levels of galectin-9 resulting in an impaired motility that can be rescued upon restoring galectin-9 levels. This data validates our results obtained using *in vitro* and *in vivo* models and highlights the importance of galectin-9 in cell migration in naturally occurring human blood DC.

**Figure 9.**
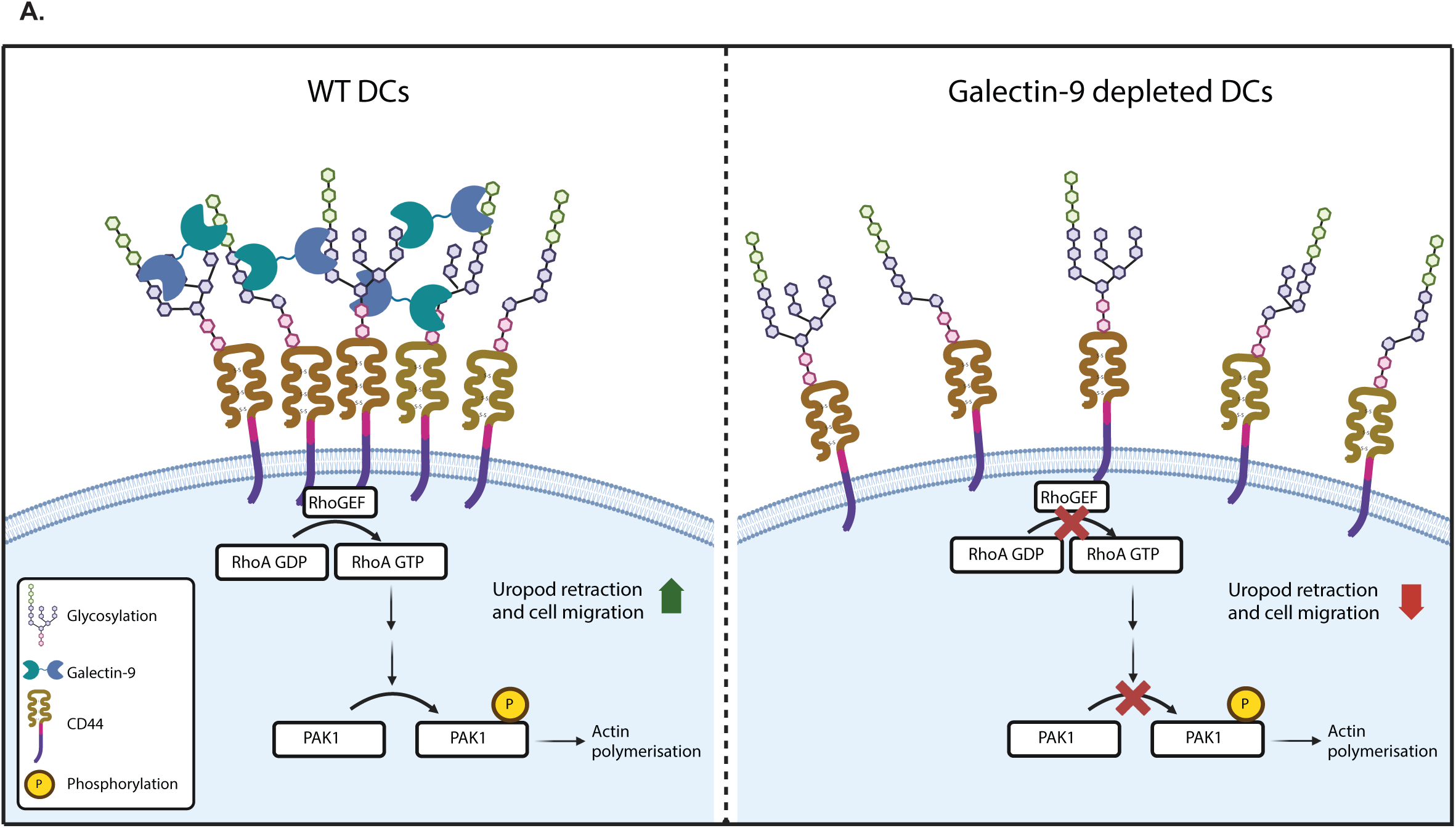
Graphical abstract. Galectin-9 controls cell rear contractility. (Left panel) Galectin-9 binds to glycosylated residues on CD44, inducing CD44 clustering and dictating the activation of RhoA downstream signalling pathways via PAK1. The activation of the galectin-9 – CD44 – RhoA axis leads to actin contractility at the cell rear and migration. In the absence of galectin-9 (right panel), CD44 clustering is deficient, which results in defective RhoA activation and impaired uropod retraction and migration.

DC trafficking from the tumour site to lymph node structures is crucial for the effective induction of anti-tumour responses (Liu et al., 2021). However, the suppressive tumour microenvironment is known to foster DC dysfunction, impeding DC motility and subsequent launching of adaptive immunity, thereby enabling tumour progression (Imai et al., 2012; Villablanca et al., 2010). How galectin-9 shapes the immune compartment within the tumour microenvironment has scarcely been addressed despite its relevant role driving tumorigenic processes (Yang et al., 2021; Zhang et al., 2020). Here, we demonstrate that exposure of cDC2s to tumour conditioned media induced galectin-9 downregulation and impaired cell migration. Treatment with galectin-9 was sufficient to restore chemokine-driven migration in immunosuppressed blood DCs which underlies the general relevance of galectin-9 in the context of multiple immunological settings.

Galectin-9 acts as an immune suppressor in B and T cells (Cao et al., 2018; Zhu et al., 2005) while we and others have reported an stimulatory function for galectin-9 in DCs. For instance, addition of recombinant galectin-9 induced the maturation of DCs (Dai et al., 2005) and we have demonstrated galectin-9 is necessary for optimal phagocytic capacity and cytokine secretion in DCs (Querol Cano et al., 2019; Santalla Mendez et al., 2023). Interestingly, those functions are mediated by the intracellular fraction of galectin-9 whereas data presented here indicates the surface-bound pool of galectin-9 is responsible for its role on cell migration. Based on our findings, we postulate that the mechanistic link between extracellular galectin-9 and DC migration relies on the formation of galectin-9-mediated functional domains at the DC membrane, enabling proper downstream RhoA signalling crucial for leukocyte propulsion. Supporting this notion, earlier reports have demonstrated galectin-9 promotes T_H_2 migration via modulation of integrin CD61 activity (Bi et al., 2011). Similarly, galectin-9 enhanced neutrophil adhesion via the regulation of integrin (CD11b and CD18) activity at the cell membrane (Iqbal et al., 2022). This is in contrast to data showing a role for intracellular galectin-3 in DC migration (Hsu et al., 2009; Kataoka et al., 2019), which highlights how different galectins exert distinct functions and that their mechanisms of action might be cell specific. Galectin-9 may function via different mechanisms and it cannot be excluded that intracellular galectin-9, which remains largely depleted after treatment with exogenous protein, is also involved in DC migration (Querol Cano et al., 2019). It is also worth mentioning that endogenous galectin-9 is highly susceptible to being cleaved in solution and therefore the used commercial recombinant galectin-9 has a shortened linker for additional stability, which may affect its binding or cross-linking abilities. Overall, data presented here indicates that galectin-9 mediated domains at the plasma membrane enable intracellular signalling resulting in cell contractility and forward movement.

Our findings establish galectin-9 as a novel regulator of both basal and chemokine-driven DC migration. DC maturation and CCR7 expression were normal in galectin-9 depleted DCs ruling out defective cell maturation as the underlying mechanism. We demonstrate that the active GTP-bound fraction of RhoA was diminished in galectin-9 KD DCs, providing the molecular basis for the observed uropod retraction defect as RhoA-mediated cytoskeletal reorganisations are indispensable for DC migration. Rho GTPase-mediated signalling is dynamically regulated by many GEFs and GAPs, which in turn form complexes with a variety of other proteins and are key factors in the spatiotemporal activation of RhoA (Lawson and Ridley, 2018). Future work could focus on studying these protein-protein interactions and comparing their levels in WT and galectin-9-depleted DCs. The most well-studied signalling pathway downstream of RhoA is that mediated by ROCK, leading to phosphorylation of the myosin light chain (MLC) and thereby increasing actomyosin contractility. RhoA can also directly bind and activate mDia to enhance actin polymerisation (Clayton and Ridley, 2020). Interestingly, no changes in either mDia expression or the phosphorylation rate of MLC were observed in galectin-9 depleted DCs despite their defect in RhoA activity. Nonetheless, the activity of other RhoA-downstream targets like Pak1 was found to correlate with galectin-9 levels, suggesting a specific regulation of the Pak pathway by galectin-9-dependent interactions. A similar phenotype has been described during contraction of smooth muscle cells in which ROCK mediated actin polymerisation via Pak phosphorylation but not MLC (Zhang et al., 2018). Pak1 is involved in cytoskeletal remodelling and is activated by phosphorylation at Thr423 mediated by Rac1, Cdc42, and RhoA (Mayhew et al., 2007; Szabo et al., 2022; Zhang et al., 2018). Decreased phosphorylation at this site indicates a lack of RhoA activity, and in general a decreased polarised state of galectin-9 KD cells. Interestingly, our analysis of publicly available datasets confirmed a correlation between RhoA-related pathways and migration of natural Killer cells is accompanied by increased RhoA transcript levels upon galectin-9 treatment, suggesting that galectin-9 control over RhoA signalling is conserved across immune cell types (Wang et al., 2016).

CD44 is located to uropods in migrating cells and can directly modulate RhoA activity *via* its cytosolic tail (Bourguignon, 2008; Bourguignon et al., 2010; Zhang et al., 2014b). Here we demonstrate that galectin-9 interacts with CD44 in DCs in line with reported interactions in neutrophils, natural Killer and T cells where it regulates cell adhesion to the CD44 substrate hyaluronan (HA) (Iqbal et al., 2022; Rahmati et al., 2023). Furthermore, our data indicates that the axis galectin-9-CD44 is also relevant in modulating cell adhesion to other substrates outside of HA. Blocking CD44 binding to galectin-9 abrogated DC migration to a greater extent in transwell chemotactic assays than in 3D collagen matrices implying that CD44 requirements for DC motility may depend on the environment or the availability of specific ligands. We hypothesise that galectin-9 acts by stabilising or anchoring CD44 in specific microdomains at the DC membrane to sustain CD44-RhoA downstream cytoskeletal reorganisations. CD44 has been shown to accumulate in lipid rafts in T cells in a process dependent on palmitoylation (Guo et al., 1994). CD44 is also highly glycosylated with O- and N-glycans and thus galectin-mediated interactions may also determine CD44 membrane localisation and its interactions with cell surface and cytoplasmic proteins. Supporting this, mutation of any of the five N-glycosylation sites of human CD44 result in the loss of CD44-mediated adhesion to HA (Bartolazzi et al., 1996). Similarly, N-glycosylation regulates CD44 interaction with Hepatic transmembrane 4 L six family member 5 (TM4SF5) and with podoplanin, required for directional migration of epithelial cells (Liao et al., 2022). CD44 becomes activated after clustering is induced (Zhang et al., 2014a) but mechanistic insights into the molecular mechanisms governing CD44 activation are largely lacking. Alternatively, galectin-mediated interactions may also determine CD44 clustering, thereby dictating receptor activation and the initiation of downstream signalling. Supporting this, inhibition of N-glycosylation reduced the average density of CD44 molecules in clusters as well as CD44 cluster size (Lakshminarayan et al., 2014) and galectin-9 binding to CD44 directly induced CD44 downstream signalling in natural Killer cells and neutrophils (Dunsmore et al., 2021; Rahmati et al., 2023).

Taken together, our study demonstrates the importance of galectin-9 in DC basal and directed migration towards lymph nodes and tumours. Furthermore, we provide for the first time evidence that galectin-9 regulates cell polarity and uropod contraction, namely by modulating RhoA activation in response to CD44 binding. Lastly, data presented here highlights a role for galectin-9 in promoting DC motility in the tumour microenvironment, underscoring galectin-9 as a target in DC-mediated anti-tumour immunity.

## Materials and Methods

### Generation of monocyte-derived dendritic cells

Dendritic cells were derived from peripheral blood monocytes isolated from a buffy coat (Sanquin, Nijmegen, The Netherlands) (de Vries et al., 2002). Monocytes isolated from healthy blood donors (informed consent obtained) were cultured for up to five days in RPMI 1640 medium (Life Technologies, Bleiswijk, Netherlands) containing 10 % foetal bovine serum (FBS, Greiner Bio-one, Alphen aan den Rijn, Netherlands), 1 mM ultra-glutamine (BioWhittaker), antibiotics (100 U/ml penicillin, 100 µg/ml streptomycin and 0.25 µg/ml amphotericin B, Life Technologies), IL-4 (500 U/ml, Miltenyi Biotec) and GM-CSF (800 U/ml, #130-093-868, MiltenyiBiotec) in a humidified, 5 % CO_2_. On day 3, medium was refreshed with new IL-4 (500 U/ml, Miltenyi Biotec) and GM-CSF (800 U/ml, Miltenyi Biotec). On day 6, moDCs were supplemented with a maturation cocktail: IL-6 (15 ng/ml, #130-093-933, Miltenyi Biotec), TNF-α (10 ng/mg, #130-094-014 Miltenyi Biotec), IL-1β (5 ng/ml, #130-093-898 Miltenyi Biotec) and PGE2 (10 ug/ml, Pfizer). When necessary, moDCs were treated with recombinant galectin-9 protein (AF2045, R&D systems) at a final concentration of 1 µg/ml.

### Isolation and culture of primary cells

Human cDC2s were isolated from peripheral blood mononuclear cells derived from healthy individuals (Sanquin, Nijmegen, the Netherlands) using the MACS CD1c+ isolation kit (130-119-475, Miltenyi Biotec) according to manufacturer’s instructions. Cell purity was determined by flow cytometry (Figure S7A) using antibodies specific against CD20-FITC (1:100 345792, BD Biosciences), CD14-PerCP (1:50, 325632, Biolegend), CD11c-APC (1:50 559877, BD Biosciences) and CD1c-PE (1:50, #130-113-302, Miletnyi Biotec). After isolation, fresh cDC2s were cultured in X-VIVO-15 (Lonza) supplemented with 2 % human serum (HS, Sigma-Aldrich) at a concentration of 0.5×10^6^ cells/ml. cDC2s were matured overnight as previously described with a maturation cocktail (MC) composed of 50 U/ml GM-CSF, 100 U/ml IL-6, 100 U/ml IL-1β, 50 U/ml TNFα (50 U/ml and 200 nM of PGE2. The next day, cDC2s were harvested, washed, and replated in media composed of 50 % fresh X-VIVO-15 supplemented with 2 % HS and 50 % melanoma cell line A375 derived conditioned media (CM) for 24 hours. When necessary, 1 µg/ml of recombinant galectin-9 was provided for the final 2 hours of cDC2 coculture with melanoma-derived CM. Next, cDC2s were harvested and washed prior to further analysis.

For BMDCs, bone marrow was taken from the tibias and femurs of 9-week-old C57BL/6 *Lgals9* ^-/-^ mice or wild type litter mates and cultured in RPMI containing 10 % FBS and 3 % granulocyte macrophage colony-stimulating factor (mGM-CSF) (Peprotech) for 7 days. Cells were treated with 1 µg/ml LPS for 16 h prior to being used. All murine studies complied with European legislation (directive 2010/63/EU of the European Commission) and were approved by local authorities (CCD, The Hague, the Netherlands) for the care and use of animals with related codes of practice.

### Cell culture, generation of tumour spheroids and conditioned media preparation

The melanoma cell lines MEL624 and BLM were cultured in Gibco DMEM high glucose medium (Life Technologies) supplemented with 10 % FBS and 0.5 % antibiotics (100 U/ml penicillin, 100 µg/ml streptomycin and 0.25 µg/ml amphotericin B, Life Technologies). A375 melanoma cell line was cultured in DMEM (high glucose, GlutaMAX, Gibco) and further supplemented with 10% FBS (HyClone) and 1% antibiotic-antimycotic (Gibco). To generate tumour spheroids cells were harvested using PBS containing 0.25 % trypsin and 4 mM EDTA and collected in Gibco DMEM high glucose medium (Life Technologies) containing 10 % FBS. For spheroid production, tumour cells were cultured in 30 µl droplets containing 4000 tumour cells resuspended in spheroid medium (60% growth medium and 40 % low viscosity methyl cellulose medium (25cp viscosity; Sigma Life Science), to which 3.6 µl of PureCol Type I Collagen (3.1 mg/ml stock solution; Advanced Biomatrix) was added.

To obtain melanoma-derived CM, A375 cells were seeded at a cell concentration of 0.25 ×10^6^ cells/ml. After 72 hours, A375 CM was harvested and centrifuged at 1500 rpms for 5 minutes to get rid of cellular debris and frozen until further use.

### Small interfering RNA knockdown

On day 3 of DC differentiation, cells were harvested and subjected to electroporation. Three custom stealth small interfering RNA (siRNA) were used to silence galectin-9 (LGALS9HSS142807, LGALS9HSS142808 and LGALS9HSS142809) (Invitrogen). Equal amounts of the siRNA ON-TARGETplus non-targeting (NT) siRNA#1 (Thermo Scientific) were used as control. Cells were washed twice in PBS and once in OptiMEM without phenol red (Invitrogen). A total of 15 μg siRNA (5 μg from each siRNA) was transferred to a 4-mm cuvette (Bio-Rad) and 5-10×10^6^ DCs were added in 200 μl OptiMEM and incubated for 3 min before being pulsed with an exponential decay pulse at 300 V, 150 mF, in a Genepulser Xcell (Bio-Rad, Veenendaal, Netherlands), as previously described (Querol Cano et al., 2019; Santalla Mendez et al., 2023). Immediately after electroporation, cells were transferred to preheated (37 °C) phenol red–free RPMI 1640 culture medium supplemented with 1 % ultraglutamine, 10 % (v/v) FCS, IL-4 (300 U/ml), and GM-CSF (450 U/ml) and seeded at a final density of 5×10^5^ cells/ml.

### Chemotaxis assays

Day 6 mature NT or *LGALS9* siRNA transfected moDCs (1×10^5^ cells in 50 µl) were seeded in the top chamber of a 24-well transwell containing a polycarbonate filter of 5 µm pore size (Corning; #CLS3421). 550 µl of RPMI medium supplemented with 1 µg/ml recombinant human CCL21 (BioLegend; #582208) or nothing as negative control were added to the lower chambers. Plates were incubated for the specified time points at 37 °C, 5 % CO_2_ after which migrated cells in the bottom chamber were collected and acquired on a MACSQuant Analyzer 10 Flow Cytometer using the MACSQuantify software (Miltenyi Biotec). The percentage of specific migration was calculated by dividing the number of cells migrated to the lower well by the total cell input (50 µl cell suspension directly measured on the MACSQuant Flow Cytometer).

Chemotactic assays using cDC2s were performed using the transwell 96-well plate 5 µm pore size (CLS3388-2EA). 3×10^4^ cDC2s were seeded in the top chamber of the transwell. The lower compartment of the transwell plate was loaded with 200 µl of X-VIVO-15 supplemented with 2 % HS and 100 ng/ml of CCL19 and CCL21 (#582102, #582202, Biolegend). To determine passive cDC2 migration, media without CCL19 nor CCL21 was used. The plate was incubated for 3 hours at 37 °C, 5 % CO_2_ after which migrated cells in the bottom chamber were collected and acquired on a MACSQuant Analyzer 10 Flow Cytometer together with the initially loaded cDC2s for each condition. Percentage of cDC2 migration was assessed by dividing the number of migrated cDC2s by the number of initially loaded cDC2s for each condition. Relative cDC2 migration was determined by dividing the percentage of migrated cDC2s of each condition by the percentage of migrated mature cDC2s unexposed to melanoma-derived CM x 100.

### 3D migration assays

A collagen mixture was generated using *PureCol*® Type I Bovine Collagen Solution (Advanced Biomatrix, final concentration 1.7 mg/ml), α-modified minimal essential medium (Sigma Albrich) and sodium bicarbonate. Collagen mixture was allowed to pre-polymerise for 5 minutes at 37 °C prior to adding a 45 µl cell suspension containing 30.000 mature day 8 NT or *LGALS9* siRNA transfected moDCs in phenol red–free RPMI 1640 culture medium supplemented with 1 % ultraglutamine and 10 % FBS. The total mixture (100 µl/well) was transferred to a 96-well black plate (Greiner Bio-One; #655090) and incubated for 45 minutes at 37 °C to allow collagen polymerisation. Afterwards, 100 µl of phenol red–free RPMI 1640 culture medium supplemented with 1 % ultraglutamine and 10 % FBS was added on top of the matrices. When appropriate, galectin-9 depleted moDCs were pre-treated with 1 µg/ml recombinant galectin-9 for 3, 24 or 48 h prior to being embedded into the collagen gel. For some experiments NT and *LGALS9* siRNA transfected moDCs were treated with 10 µg/ml of the anti-CD44 blocking antibody (Iqbal et al., 2022) or rat IgG2a isotype for 30 min prior to adding recombinant galectin-9 for another 30 min. Cells were subsequently harvested and washed before being embedded into the collagen gel. When relevant, moDC cell suspension was mixed with one tumour spheroid prior to being added to the collagen matrix and the microscopy plate was inverted after every 7-10 minutes during collagen polymerisation to prevent the spheroid from sinking into the matrix. In collagen gels containing tumour spheroid moDCs were stained with PKH-26 (Sigma Albrich) or CellTrace™ CFSE or CellTrace™ Far Red (Invitrogen) according to manufacturer’s instructions and to distinguish them from tumour cells. After one day of incubation with spheroids, collagen gels were first fixed with 2 % PFA in PBS for 5 min, then 4 % PFA in PBS for 20 min, both at 37° C. To localise the spheroid, collagen gels were subsequently incubated with Alexa Fluor™ 488 phalloidin (A12379, Invitrogen) or phalloidin-iFluor 647 (ab176759, Abcam) in PBS + 3 % bovine serum albumin, 0.1 M glycine, and 0.3 % triton for 3 hours at room temperature. Imaging was performed with a Zeiss LSM880 confocal microscope, using a 10x 0.45 NA air objective (Zeiss) to make z-stacks with 5 μm intervals. To quantify spheroid infiltration and the number of moDCs surrounding the spheroid, one plane was taken about 50 μm into the spheroid and moDC count was performed by first setting a threshold and then by the analyze particles feature in Fiji (ImageJ). For moDCs surrounding the spheroid, a circular band with a fixed surface area surrounding the spheroid was taken for analysis.

Time-lapsed video microscopy was performed using the BD Pathway 855 spinning disk confocal microscope (BD Bioscience), the atto vision software (BD Bioscience) and the 10X objective (Olympus). Sequential images were acquired every 4-5 minutes for 10 hours using the 548/20 excitation filter (Chroma), emission filter 84101 (Chroma) and dichroic filter 84000 (Chroma). When appropriate, time-lapse microscopy was performed using the Celldiscoverer7 (Zeiss), using the 5x objective with 2x tube lens or the the BD Pathway 855 spinning disk confocal microscope (BD Bioscience), the atto vision software (BD Bioscience) and the 10X objective (Olympus). Sequential images were acquired every 4-5 minutes for 10 hours using the 548/20 excitation filter (Chroma), emission filter 84101 (Chroma) and dichroic filter 84000 (Chroma). Time-lapse sequences were analysed with the ImageJ manual-tracking plugin to measure cell velocity and to track individual cells.

### In vivo adoptive transfer

Wild type and galectin-9^-/-^ DCs were labelled with 5 µM carboxyfluorescein succinimidyl exter (CFSE) violet and far-red dyes following manufacturer’s instructions (Invitrogen; #C34571 and C34572 respectively), mixed in equal numbers (1×10^6^ each in 50 µl) and co-injected into the same footpad or tail vein of either wild type or galectin-9^-/-^ recipient mice. Donor DCs arriving in the draining lymph node (popliteal and inguinal respectively) were enumerated 48 h later by flow cytometry using a BD FACSLyric flow cytometer (BD BioSciences).

### Reverse phase protein array (RPPA)

Cellular proteins were denatured in a 1 % SDS + 2-mercaptoethanol buffer solution and diluted in five 2-fold serial dilutions in dilution lysis buffer. Serially diluted lysates were arrayed on nitrocellulose-coated slides (Grace Bio-Labs) by the Quanterix (Aushon) 2470 Arrayer (Quanterix Corporation) and each slide was probed with a validated primary antibody plus a biotin-conjugated secondary antibody (https://www.mdanderson.org/research/research-resources/core-facilities/functional-proteomics-rppacore/antibody-information-and-protocols.html). Signal detection was amplified using an Agilent GenPoint staining platform (Agilent Technologies) and visualized by DAB colorimetric reaction. The slides were scanned (Huron TissueScope, Huron Digital Pathology) and quantified using customized software (Array-Pro Analyzer, Media Cybernetics) to generate spot intensity. Relative protein level for each sample was determined by RPPA SPACE (Shehwana et al., 2022) (developed by MD Anderson Department of Bioinformatics and Computational Biology, https://bioinformatics.mdanderson.org/public-software/rppaspace/). The protein concentrations of each set of slides were then normalized for protein loading. Correction factor was calculated by (1) median-centring across samples of all antibody experiments; and (2) median centring across antibodies for each sample. Results were then normalized across RPPA sets by replicates-based normalization as described (Akbani et al., 2014). Details of the RPPA platform as performed by the RPPA Core are described in (Siwak et al., 2019). Pathway enrichment analysis was performed using the metascape platform and the analysis outlined in (Zhou et al., 2019). **Flow cytometry**

To determine depletion of galectin-9 following siRNA transfection, single cell suspensions were stained with a goat anti-galectin-9 antibody (AF2045, R&D systems) at 8 μg/ml or isotype control as negative control for 30 min at 4 °C. Before staining, moDCs were incubated with 2 % human serum for 10 min on ice to block non-specific interaction of the antibodies with Fc receptors. A donkey-anti goat secondary antibody conjugated to Alexa Fluor 488 was used (Invitrogen; 1:400 (v/v). moDCs were incubated for 30 min on ice with antibodies against HLA-DR (BD BioSciences; # 555811, clone G46-6, FITC-labelled), CD80 (BD BioSciences; # 557227, clone C3H, PE-labelled), CD86 (BD BioSciences; #555658, clone 2331, PE-labelled), CD83 (Miltenyi; # 130-094-186, clone HB15, APC-labelled), CCR7 (#130-094-286, Miltenyi Biotec). All antibodies were used at a final 1:25 (v/v) dilution in cold PBS containing 0.1 % BSA, 0.01% NaH_3_ (PBA) supplemented with2 % HS.

To phenotype cDC2s, harvested cDC2s were first blocked in PBA buffer supplemented with 2 % HS for 15 minutes. After blocking, cDC2s were stained in PBA buffer with anti-galectin-9 antibody (1:50, AF2045, R&D systems) and anti-CCR7 (1:200) for 20 minutes. Next, cells were washed and stained with a secondary antibody, a donkey-anti-goat antibody conjugated to Alexa488 (1:400, Invitrogen) for 20 minutes. Stained cDC2s were analysed by flow cytometry using a FACS Verse (BD, Frankllin lakes, NJ, USA) and later analysed using FlowJo software (BD, Franklin lakes, NJ, USA).

### RhoA pull down

2–3×10^6^ day 6 WT, galectin-9 KD or galectin-9 rDCs were collected. RhoA GTPase activity was measured using the RhoA Pull-Down Activation Assay Biochem Kit (#BK036; Cytoskeleton) according to the manufacturer’s instructions. Active RhoA protein was quantified using the Image Studio Lite software (Li-Cor).

### Western Blot

Day 6 or 7 moDCs were lysed in lysis buffer for 30 min on ice prior to being spun down at 10.000 rpm for 5 min. The BCA protein assay (Pierce, ThermoFisher scientific) was conducted to determine protein concentration and following manufacturer’s instructions and for each sample, 20 µg of total protein were diluted using SDS sample buffer (62.5 mM Tris pH 6.8, 2 % SDS, 10 % glycerol).

Proteins were separated by SDS-PAGE and blotted onto PVDF membranes. Membranes were blocked in TBS containing 3 % BSA at room temperature for 1 h prior to be stained with specific antibodies. Antibody signals were detected with fluorophore coupled secondary antibodies and developed using Odyssey CLx (Li-Cor) following manufacturer’s instructions. Images were retrieved using the Image Studio Lite 5.0 software. The following primary antibodies were used for Western Blotting: goat anti-Galectin-9 (AF2045, R&D systems, Minneapolis, Minnesota) at 1:1000 (v/v), rabbit anti-phospho(Thr 423) Pak1 (#2601; Cell Signaling Technology) at 1:500 (v/v), rabbit anti-Pak1 (#2602; Cell Signaling Technology) at 1:500 (v/v), rabbit anti-GAPDH (#2118; Cell Signaling Technology) at 1:500 (v/v), mouse anti-phospho myosin light chain (#3675, Cell Signaling Technology) at 1:500 (v/v), mouse anti-mDia (#610848, BD BioSciences) at 1:500 (v/v) and rat anti-tubulin (Novus Biological, Abingdon, United Kingdom) at 1:2000 (v/v). The following secondary antibodies were used: donkey anti-goat IRDye 680 (920-32224, Li-Cor, Lincoln, Nebraska), donkey anti-rabbit IRDye 800 (926-32213, Li-Cor), donkey anti rabbit IRDye 680 (926-68073, Li-Cor), goat anti rabbit IRDye 800 (926-32211, LiCor), goat anti-rat IRDye 680 (A21096, Invitrogen, Landsmeer, Netherlands), donkey anti-mouse IRDye 680 (926-68072, Li-Cor). All secondary antibodies were used at 1:5000 (v/v).

### Data analysis

Cell tracking was performed using the manual tracking plugin of Fiji (ImageJ) with adjusted microscope-specific time and calibration parameters. Individual cells were tracked for at least 90 minutes 5 hours after being embedded in the collagen to allow adaptation to the environment. The mean square displacement (MSD) over time intervals was determined as previously described (van Rijn et al., 2016). In short, the MSD was calculated per time interval for each cell. The average per time interval was calculated for all cells corrected for the tracking length of the cells. The Euclidean distance reached by a cell after 60 minutes of tracking with respect to their starting position was calculated using the Chemotaxis and Migration software (Ibidi) after adjusting the X/Y calibration and time acquisition interval. Tracking plots were generated using the Chemotaxis and Migration software (Ibidi).

For the pathway enrichment analysis, we identified the top 50 genes correlating with *lgals9* gene across all immune cells using the ULI RNA-seq dataset (GSE109125) and the gene constellation tool from the Immunological Genome Project (ImmGen). We then conducted a functional pathway enrichment analysis using gProfiler (Kolberg et al., 2023) accross the Reactome dataset (Griss et al., 2020).

All data was processed using Excel 2019 (Microsoft) and plotted using GraphPad Prism 8 software. All statistical analysis was done using Prism 8. All data is expressed as mean ± SEM unless otherwise stated. The statistical test used to analyse each data set is described in the corresponding figure legend. Unpaired t-test was used to compare mean values between NT and *gal9* siRNA transfected cells. One-way analysis of variance (ANOVA) was used for multiple comparison. cDC2 data was analysed using one-way analysis of variance (ANOVA) followed by a Dunnett’s test for multiple comparison. Statistical significance was considered for p values < 0.05.

## Acknowledgements

The authors thank the Radboudumc Technology Center Microscopy for use of their microscopy facilities and Prof. Geert van den Bogaart for his help in analysing individual cell tracking. The RPPA Core is supported by NCI Grant # CA016672 and Dr. Yiling Lu’s NIH R50 Grant # R50CA221675: Functional Proteomics by Reverse Phase Protein Array in Cancer. “The Functional Proteomics Reverse Phase Protein Array Core was supported in part by The University of Texas MD Anderson Cancer Center, P30CA016672, and R50CA221675.” This work is supported by a PhD grant from Radboudumc, grants 11618 and 12949 from the Dutch Cancer Society. AvS is supported by the Netherlands Organization for Scientific Research (NWO): the Institute of Chemical Immunology (project ICI 000-23), ZonMW (project 09120012010023), and the European Research Council: Consolidator Grant (project 724281) and Proof-of-Concept Grant (project 101112687). The authors declare no competing financial interests.

---

**Supplementary Figure 1.**
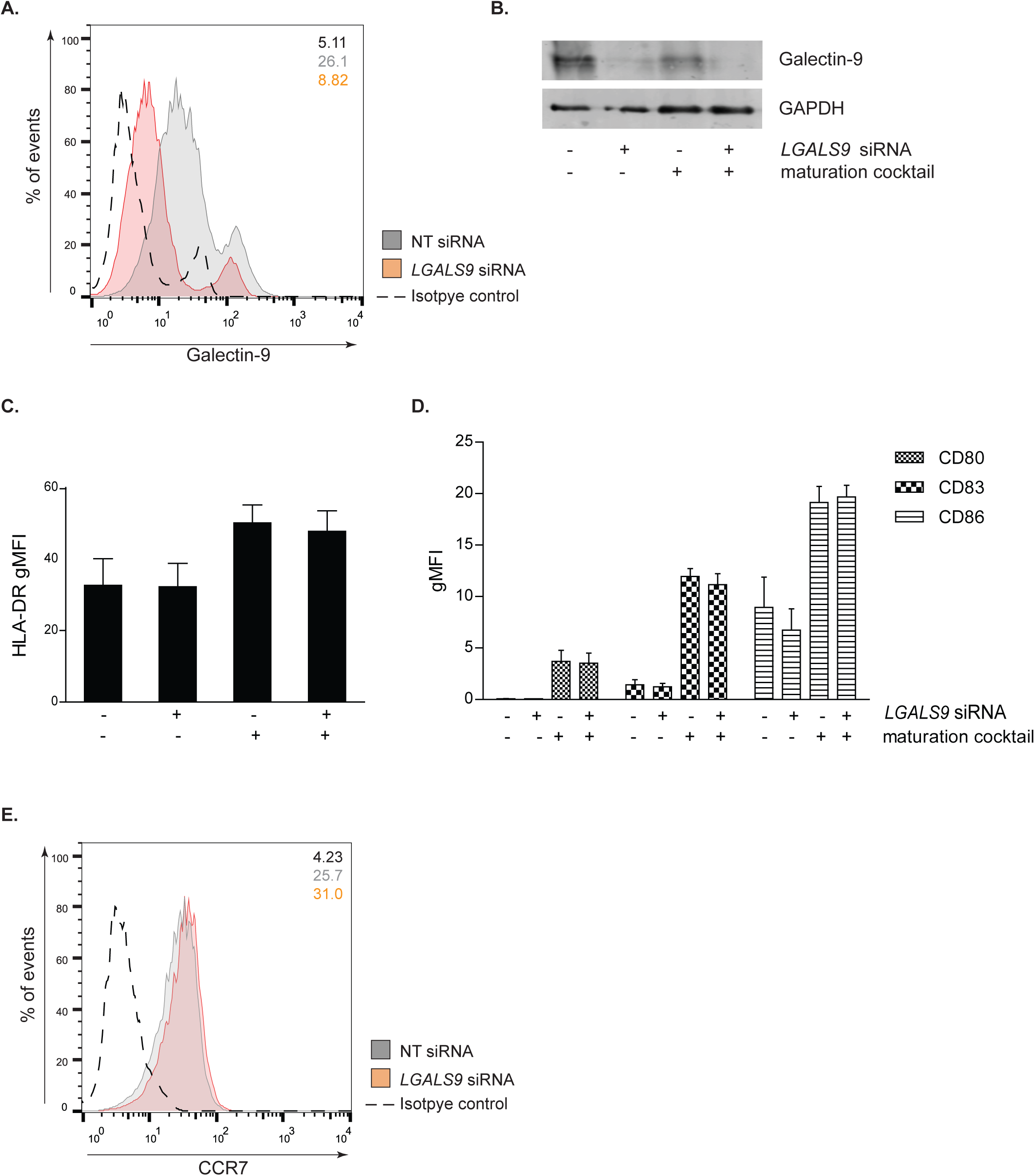
Galectin-9 depletion does not affect dendritic cell maturation. **A.** Day 3 moDCs were transfected with *LGALS9* siRNA or a NT siRNA. Levels of galectin-9 were assessed by flow cytometry 48 h after transfection. Flow cytometry graph shows galectin-9 levels at the plasma membrane. Graph depicts a representative donor and numbers in inset indicate geometric mean fluorescence intensity (gMFI). NT siRNA (light grey population), *LGALS9* siRNA-transfected moDCs (red population). Black dotted line represents isotype control values. **B.** Total lysates from NT and *LGALS9* siRNA transfected cells treated with maturation cocktail or nothing were subjected to Western Blot and galectin-9 expression analysed. GAPDH was used as loading control. Immunoblot is representative of four independent experiments. **C** and **D.** NT or *LGALS9* siRNA transfected moDCs were matured and levels of HLA-DR, CD80, CD86 and CD83 analysed by flow cytometry. Graph depicts mean ± SEM surface protein levels (gMFI) of four independent donors. **E.** Membrane levels of CCR7 were assessed by flow cytometry in moDCs treated as per (C). Graph is a representative donor out of three analysed and numbers in inset indicate geometric mean fluorescence intensity (gMFI). NT siRNA (grey population), *LGALS9* siRNA-transfected moDCs (red population). Black dotted line represents isotype control values.

**Supplementary Figure 2.**
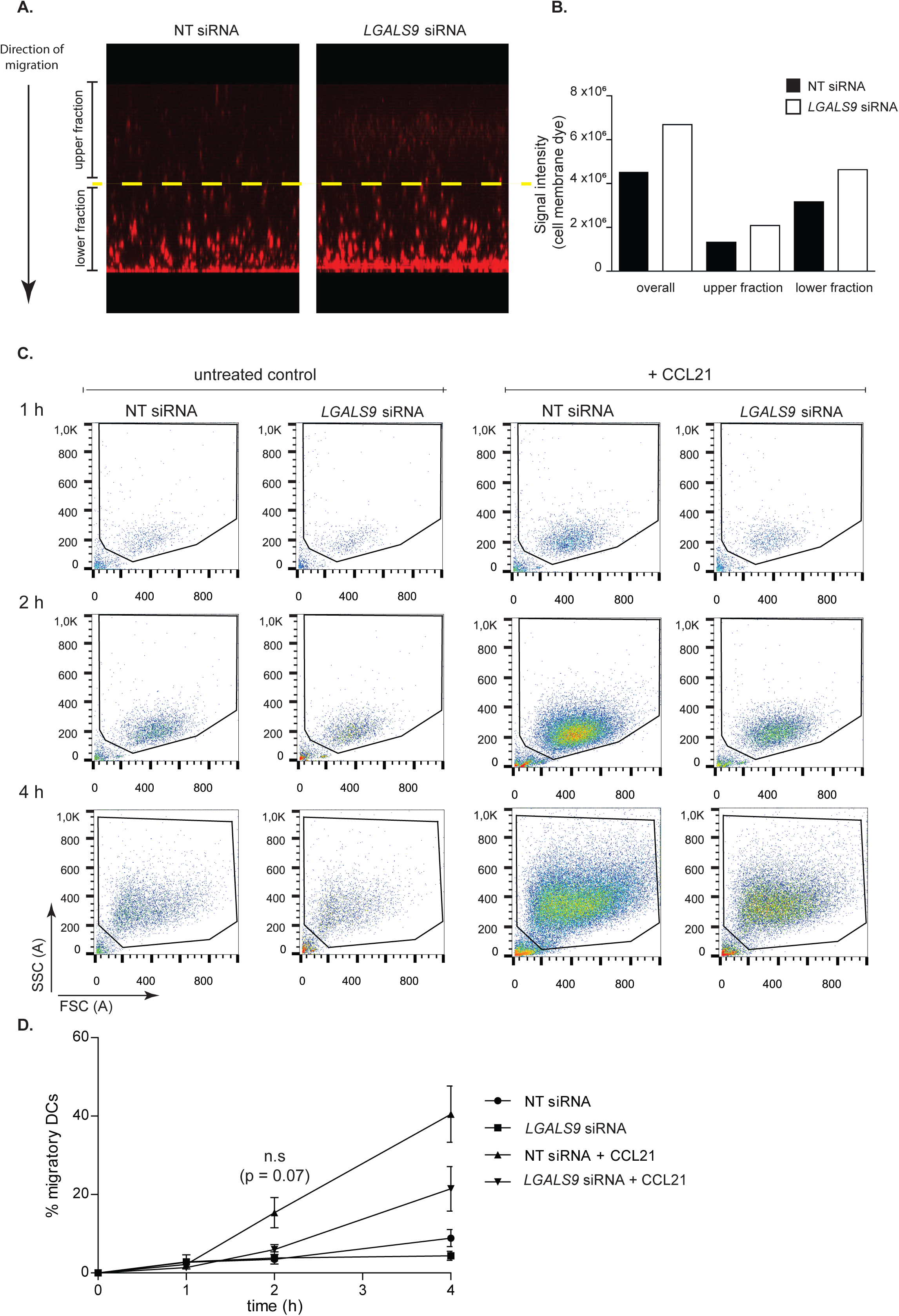
Galectin-9 depletion impairs chemokine-driven migration in dendritic cells. **A.** Mature *LGALS9* or NT siRNA transfected moDCs were stained with PKH26, seeded on top of a collagen layer overlaying a transwell chamber and subjected to a CCL21 chemokine gradient. Cells were left to migrate for 24 h after which collagen gels were fixed and imaged by confocal microscopy. **B.** Quantification of PKH signal in WT and galectin-9 KD DC containing collagens for the total length of the collagen gel as well as the upper and lower fraction. **C.** moDCs treated as per A were placed in the upper chamber of a transwell system and subjected to a CCL21 chemokine gradient. Migrated cells were collected at the bottom chamber 1, 2 and 4 h after seeding and measured by flow cytometry. Graphs show representative donor of 4 independent donors analysed. **D.** Quantification of data shown in (C). Graph shows mean percentage ± SEM percentage of migratory DCs relative to total number of seeded cells for each time point and genotype.

**Supplementary Figure 3.**
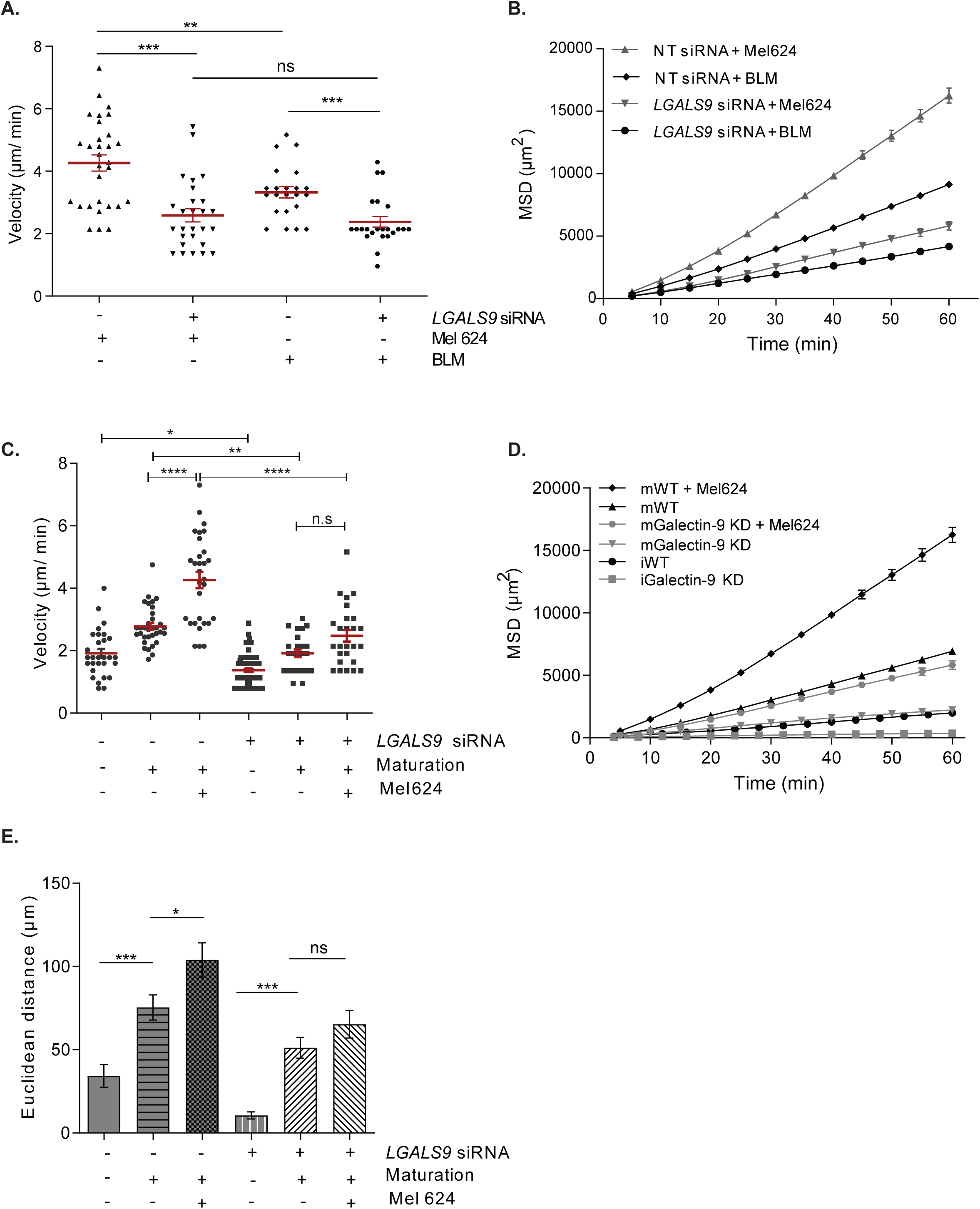
Galectin-9 depletion diminishes dendritic cell 3D migration. **A** and **B.** WT or galectin-9 depleted moDCs were matured and embedded within a 3D collagen matrix containing either a MEL624 or a BLM melanoma cell spheroid. The median cell velocity (A) and the mean square displacement (MSD) (B) were calculated. Data represents the mean value ± SEM of a representative donor. At least twenty cells were analysed per condition and condition. **C – E.** moDCs were transfected with either a NT or a *Lgals9* siRNA, matured or not and embedded in a collagen matrix containing a MEL624 tumour spheroid. The migration pattern of either immature (at day 6 of DC differentiation), mature (at day 8) and mature cells in the presence of a tumour spheroid (at day 8) was analysed for one representative donor. The median cell velocity (C), the MSD (D) and the Euclidean distance reached after 60 minutes of tracking (E) were assessed. In D: m = mature DCs; i = immature DCs. Graphs represent the mean value ± SEM of one representative donor. At least twenty-five cells were analysed per condition. One-way ANOVA was performed to compare NT and *LGALS9* siRNA transfected moDCs. n.s. p > 0.05; * p < 0.05; ** p< 0.01; *** p< 0.001; **** p< 0.0001.

**Supplementary Figure 4.**
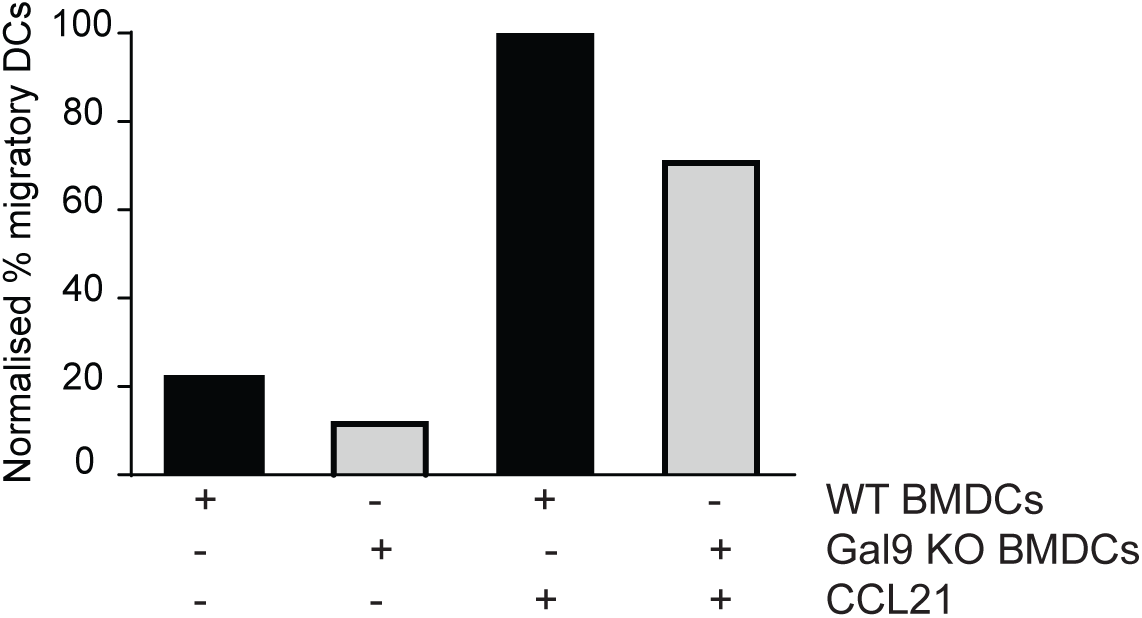
Chemokine-driven migration is impaired in murine galectin-9 KO DCs. Bone marrow derived dendritic cells (BMDCs) obtained from either wild type (WT) or galectin-null (galectin-9 ^-/-^) mice were subjected to a chemokine transwell assay in the presence or absence of chemokine CCL21. Data shows the percentage of moDCs that migrated relative to input.

**Supplementary Figure 5.**
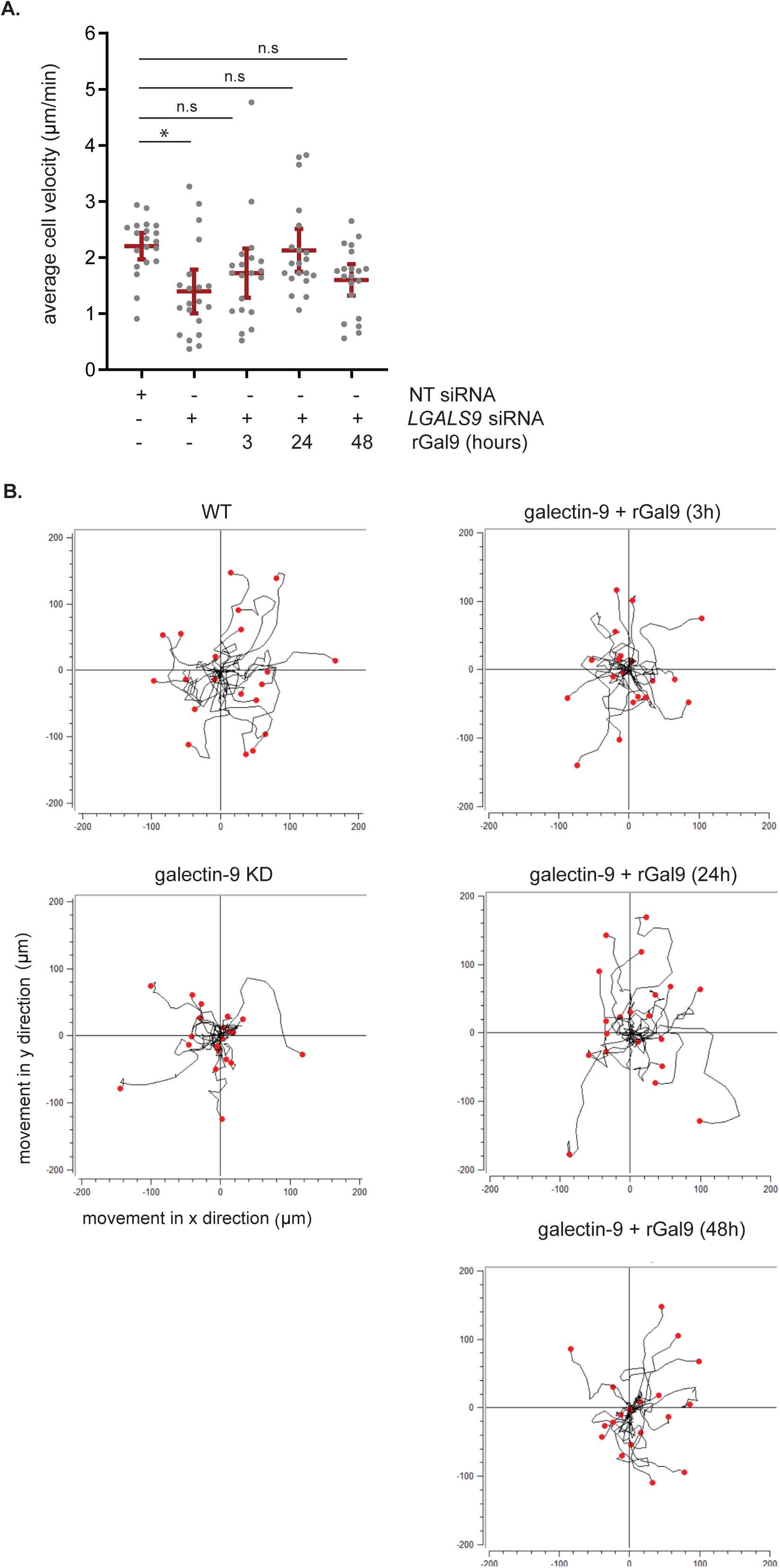
Treatment with recombinant galectin-9 protein restores dendritic cell migration. **A.** WT, galectin-9 KD and galectin-9 KD moDCs treated with 1 µg/ml recombinant galectin-9 protein (gal-9 KD + rGal9 moDCs) for either 3, 24 or 48 h were embedded in a 3D collagen matrix and their velocity calculated. Graph depicts average velocity ± SEM for one representative donor out of 4 analysed. **B.** Individual trajectory plots of cells shown in (A). End points of tracks are indicated by red dots. One way ANOVA with a Kruskal-Wallis for multiple comparison test was performed. n.s. p > 0.05; * p < 0.05; *** p< 0.001.

**Supplementary Figure 6.**
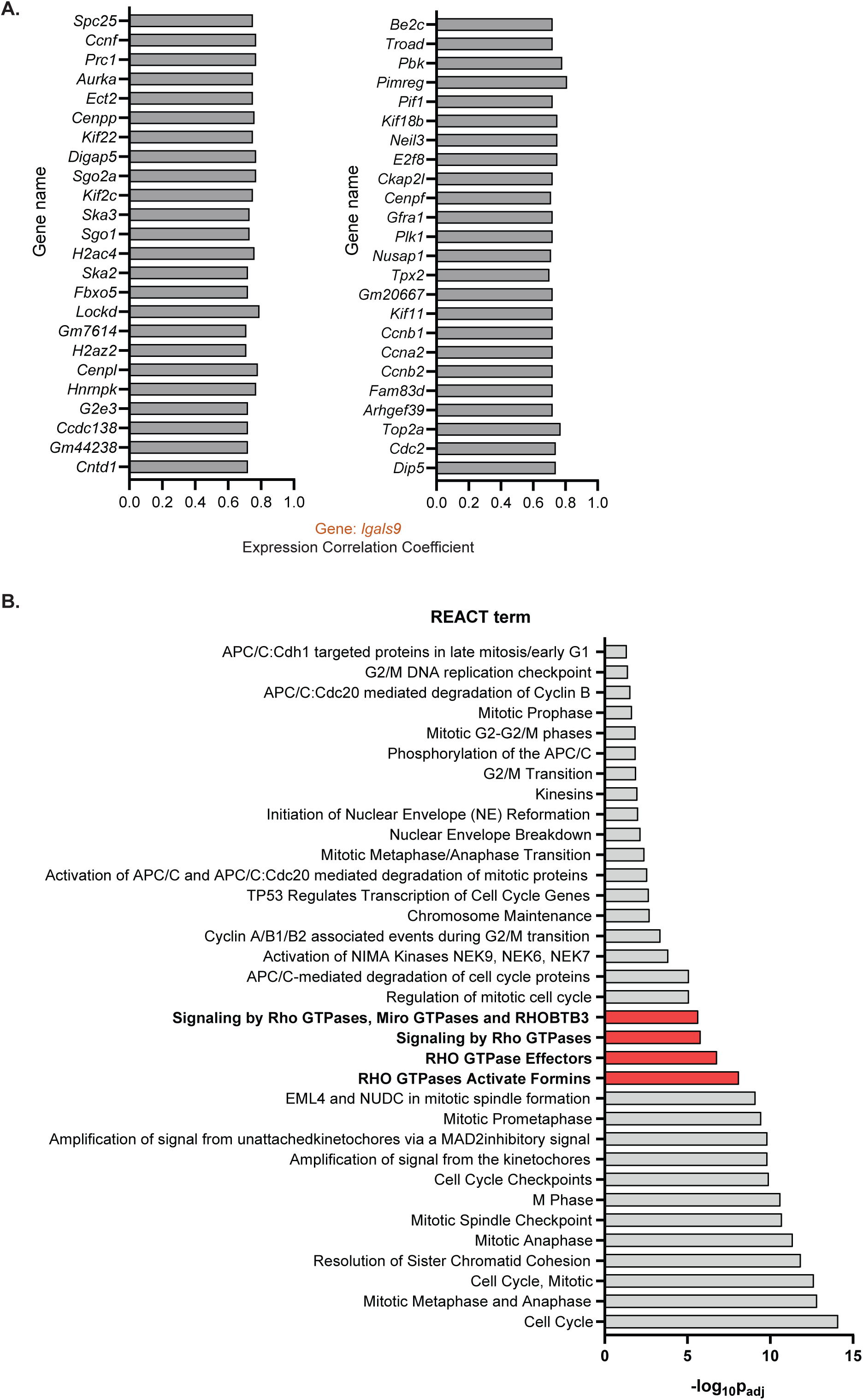
RhoGTPase-mediated pathways positively correlate with *lgals9* expression. **A.** 50 genes that positively correlated to *lgals9* gene across all immune cell types (GSE109125 dataset) grouped by secondary correlation. **B.** Functional pathway enrichment analysis (Reactome dataset). Data is shown as -log_10_ of the adjusted p-value.

**Supplementary Figure 7.**
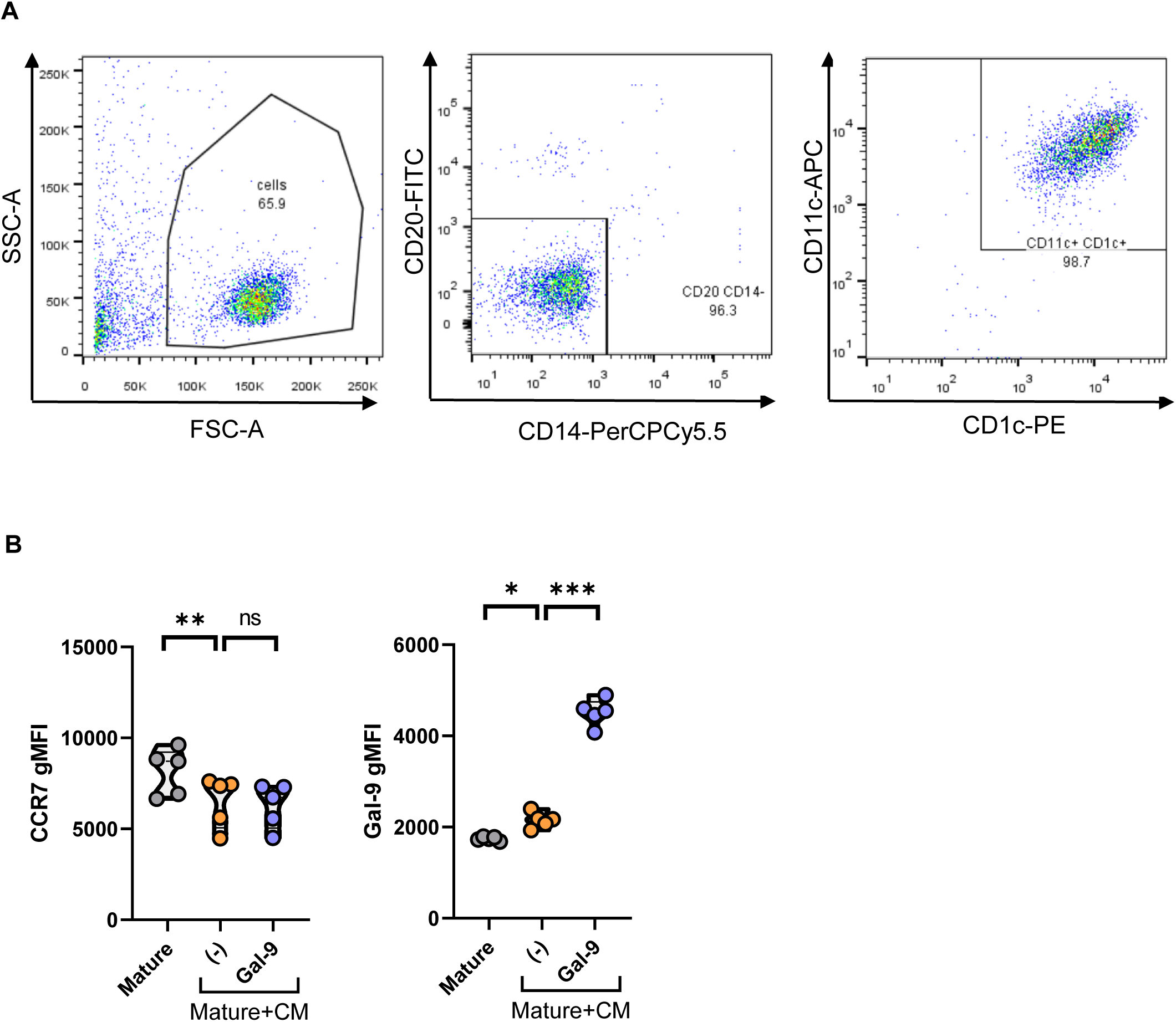
cDC2 cell purity and supplementary phenotype analysis. **A.** Gating strategy to determine cDC2 purity after isolation from PBMCs. cDC2s were stained for CD1c, CD11c, CD14 and CD20. A first gate is set based on the physical FCS-A/SSC-A parameters followed by selecting on negative cells for CD14 and CD19. cDC2s are then identified as positive cells for CD11c and CD1c. **B.** Surface galectin-9 and CCR7 expression in galectin-9 and CCR7 positive cDC2s. Graph shows average ± SEM geometric mean fluorescence intensity (gMFI) of 5 individual donors. One-way ANOVA followed by Dunnett’s test for multiple comparison was performed. * p < 0.05; ** p < 0.01; *** p < 0.001.

